# *Scribble* mutation disrupts convergent extension and apical constriction during mammalian neural tube closure

**DOI:** 10.1101/2020.09.18.303446

**Authors:** Alyssa C. Lesko, Raymond Keller, Ping Chen, Ann Sutherland

## Abstract

Morphogenesis of the vertebrate neural tube occurs by elongation and bending of the neural plate, tissue shape changes that are driven at the cellular level by polarized cell intercalation and cell shape changes, notably apical constriction and cell wedging. Coordinated cell intercalation, apical constriction, and wedging undoubtedly require complex underlying cytoskeletal dynamics and remodeling of adhesions. Mutations of the gene encoding Scribble result in neural tube defects in mice, however the cellular and molecular mechanisms by which Scrib regulates neural cell behavior remain unknown. Analysis of Scribble mutants revealed defects in neural tissue shape changes, and live cell imaging of mouse embryos showed that the Scrib mutation results in defects in polarized cell intercalation, particularly in rosette resolution, and failure of both cell apical constriction and cell wedging. *Scrib* mutant embryos displayed aberrant expression of the junctional proteins ZO-1, Par3, Par6, E- and N-cadherins, and the cytoskeletal proteins actin and myosin. These findings show that Scribble has a central role in organizing the molecular complexes regulating the morphomechanical neural cell behaviors underlying vertebrate neurulation, and they advance our understanding of the molecular mechanisms involved in mammalian neural tube closure.

**Highlights:**

- Polarized cell intercalation is lost in *Scrib* mutant embryos
- *Scrib* mutation has specific effects on rosette formation and resolution
- *Scrib* mutation disrupts apical constriction and cell shape changes necessary for neural tube closure
- Adherens and tight junction composition is altered in the neural epithelial cells of *Scrib* mutants

## Introduction

Neural tube development in vertebrates begins with the elongation of the neural plate, followed by the bending, folding, and fusion of the plate into a tube (neural tube closure, NTC) (Nikolopoulou et al., 2017; Sadler, 2005). In mice, a “zippering” mechanism is initiated sequentially at three sites, at closure 1 at the hindbrain/cervical boundary, at closure 2 at the forebrain/midbrain boundary, and at closure 3 at the most rostral point of the forebrain, and it proceeds in both directions from these sites. (Copp and Greene, 2010; Nikolopoulou et al., 2017; Sadler, 2005). Failure of any of the closures to form, or of subsequent zippering, results in severe birth defects depending on where the failure occurs (Copp and Greene, 2013; Greene and Copp, 2014). For example, failure of zippering between closures 1 and 2 or between closures 2 and 3 result in exencephaly or anencephaly (Copp and Greene, 2013) whereas failure of zippering in the caudal spinal region results in spina bifida. Failure of closure 1 to form, and subsequently failure of zippering, results in an open hindbrain and spinal cord and is classified as craniorachischisis (CRN) (Copp and Greene, 2013). All of these neural tube defects (NTD) are detrimental to quality of life and often result in death at or before birth.

Elongation of the neural plate during NTC is driven by convergent extension (CE), a process in which cells intercalate mediolaterally (ML), thereby forming a narrower, longer array (Keller, 2002). In mesenchymal tissues, cells intercalate by extending mediolaterally oriented protrusions between one another, attach, and shorten to crawl between neighboring cells, thus elongating the tissue in the anterior-posterior dimension (Keller, 2002; Munro and Odell, 2002). Examples of this are the mesodermal cells of *Xenopus* (Shih and Keller, 1992), mouse somitic mesodermal cells (Yen et al., 2009), and ascidian notochordal cells (Munro and Odell, 2002). Epithelial cells can also rearrange, despite their apical junctional complexes. This was initially demonstrated during eversion of imaginal discs (Fristrom, 1976; Taylor and Adler, 2008), during germband extension in *Drosophila* (Irvine and Wieschaus, 1994), and in the *Xenopus* marginal zone (Keller, 1978). Analysis of germ band extension in *Drosophila* showed that rearrangement of apical intercellular junctions occurs by remodeling through a “T1” process or through rosette formation and resolution (Bertet et al., 2004; Blankenship et al., 2006). However, epithelial cells also use basolateral cell protrusive activity to rearrange. The dorsal epithelial cells of *C. elegans* embryos (Williams-Masson et al., 1998) extend large basolateral protrusions between neighboring cells, followed by apical intercalation. Similarly, mouse neural plate epithelial cells also intercalate by extending mediolaterally polarized protrusions from their basolateral ends in a crawling mechanism not unlike mesenchymal cells (Williams et al., 2014). Later work in Drosophila confirmed that germ band elongation is also accompanied by a combination of basolateral protrusive activity and apical junctional rearrangement (Sun and Amourda, 2017).

The bending and closure of the neural epithelium is driven primarily by cell shape changes: apical constriction, apical basal elongation, and cell wedging (reviewed in (Nikolopoulou et al., 2017; Sutherland et al., 2020)). As the tissue elongates anterior-posteriorly, cells in the neural plate elongate and orient their basal ends mediolaterally and become increasingly columnar while constricting their apical membrane to form wedge shaped cells (Moury and Schoenwolf, 1995; Suzuki et al., 2012). These cell shape changes provide the mechanical force necessary for the bending and zippering of the neural plate (Inoue et al., 2016; Nikolopoulou et al., 2017).

The planar cell polarity (PCP) pathway, which regulates the planar orientation of cells in a tissue, has been shown to affect NTC through orientation of cell intercalation and promotion of apical constriction (Axelrod and McNeill, 2002; Heisenberg et al., 2000; López-Escobar et al., 2018; Tada and Smith, 2000; Wallingford and Harland, 2001; Wallingford et al., 2000; Williams et al., 2014; Ybot-Gonzalez et al., 2007). Mutation of genes encoding PCP components such as Frizzled (Fz3/Fz6 or Fz2/7), Van Gogh-like (Vangl1/Vangl2), and Dishevelled (Dvl1/2/3) in mice cause open neural tube and CRN (Hamblet et al., 2002; Kibar et al., 2001; Murdoch et al., 2001a; Wang et al., 2006a; Wang et al., 2006b). *Vangl2 Looptail* (*Vang2l^Lp^*) mice, which contain a point mutation in the *Vangl2* gene that prevents delivery of Vangl2 to the plasma membrane (Kibar et al., 2001; Merte et al., 2010; Murdoch et al., 2001a), and truncation mutants of the Protein tyrosine kinase 7 (*Ptk7)* gene exhibit disrupted CE and NTC (Williams et al., 2014).

The PCP pathway regulates morphogenesis and NTC by regulating cytoskeletal dynamics through Rho kinase signaling (reviewed in (Coravos et al., 2017; Heer and Martin, 2017; Sutherland and Lesko, 2020)). In chick embryos, actin, myosin II, and Rho accumulate at apical ends of cells prior to neural plate bending, and inhibition of actin polymerization and myosin result in failure of NTC (Kinoshita et al., 2008). Similarly, in the mouse neural plate inhibition of Rho kinase and actin disassembly inhibits NTC through effects on cell shape and apical constriction (Butler et al., 2019; Williams et al., 2014; Ybot-Gonzalez et al., 2007), and Rho kinase inhibition in *Vang2l^Lp+/-^* embryos exacerbated NTD (Escuin et al., 2015; Ybot-Gonzalez et al., 2007). Furthermore, mutation of *Ptk7* disrupts the localization of myosin IIB in mouse neural cells (Williams et al., 2014), and affects signaling to Rho kinase (Andreeva et al., 2014). The PCP pathway thus regulates cytoskeletal dynamics and Rho signaling to promote NTC through junctional rearrangement, cell shape changes, and biomechanical accommodation of neural fold zippering (Galea et al., 2017; Galea et al., 2018).

The protein Scribble (Scrib) acts as a scaffold to promote formation of protein complexes at the plasma membrane which are involved, particularly in epithelia, in apical-basal and planar polarization of the cell (Bonello and Peifer, 2018), and mutations in *Scrib* result in CRN in both mice and humans (Kharfallah et al., 2017; Lei et al., 2013; Murdoch et al., 2003; Robinson et al., 2012; Zarbalis et al., 2004). Scrib contains an amino-terminal LRR region important for its correct localization to the plasma membrane and for its role in regulating apical-basal polarity and junctional composition (Bonello and Peifer, 2018; Kallay et al., 2006). At the carboxy terminus, Scrib has four PDZ binding domains important for protein-protein interactions including domains that directly interact with Vangl2 and the PCP pathway (Kallay et al., 2006). Scrib’s role in neural tube development has been predominantly studied in two mouse models: *Scribble Circletail* (*Scrib^Crc^*) (Murdoch et al., 2003) and *Scribble Rumplestiltzchen (Scrib^rumz^,* also published as Line 90) (Dow et al., 2007; Zarbalis et al., 2004), both of which display CRN and short body axis phenotypes. The *Scrib^Crc^* mutation is a frameshift mutation in the PDZ domain causing the truncation of Scrib and loss of function and expression (Murdoch et al., 2003). The *Scrib^rumz^* mutant contains a point mutation in the N-terminal LRR domain that leads to decreased expression and membrane localization (Zarbalis et al., 2004). Both *Scrib^Crc^* and *Scrib^rumz^* mutants genetically interact with the *Vangl^Lp^* mutant to cause CRN (Murdoch et al., 2001b; Zarbalis et al., 2004) though the mechanisms by which Scrib and Vangl2 cooperate are not fully understood.

It is clear that the PCP pathway and Scrib influence neural tube development, however the mechanisms by which Scrib regulates NTC remain unknown. Here we show that the Scrib mutation leads to defects in both of the major, formative events in NTC: CE and apical constriction/cell wedging. We also show the it does so by altering the molecular composition of the apical junctional complex and the cytoskeleton, which results in specific defects in the cellular behaviors underlying NTC.

## Materials and Methods

### Animals

*Scrib^rumz^* (also known as *Scrib^L90^*) mice (Zarbalis et al., 2004), and *Scrib^Crc^* mice (Murdoch et al., 2003), were obtained from Dr. Xiaowei Lu, University of Virginia, and kept as heterozygous stocks in our colony. The *Scrib^rumz^* mutant contains a point mutation in the N-terminal LRR domain (Zarbalis et al., 2004) that affects both the stability of the protein and its ability to localize to the membrane (Dow et al., 2007), while the *Scrib^Crc^* mutation is a loss of function frameshift mutation in the Scrib C-terminus PDZ binding domains causing loss of Scrib expression (Murdoch et al., 2003). *Vangl2^Lp^* mice, exhibiting a point mutation that causes Vangl2 to be sequestered in the endoplasmic reticulum and not properly targeted to the membrane, were obtained from The Jackson Lab and kept as heterozygous stocks in our colony (Kibar et al., 2001; Merte et al., 2010). *Vangl1^co/co^* and *Vangl2^co/co^* conditional knockout mice were obtained from Dr. Jeremy Nathans, Johns Hopkins University and Dr. Yingzi Yang, Harvard University, respectively (Chang et al., 2016; Song et al., 2010), and kept as homozygotes in our colony. *Tg(Sox2-cre)1Amc/J (Sox2Cre*) females, obtained from The Jackson Lab, were crossed to *Vangl1^co/co^* and *Vangl2^co/co^* conditional males to produce *Vangl1^KO/+^* and *Vangl2^KO/+^* heterozygotes. *Vangl2^GFP/GFP^* mice, expressing GFP labelled Vangl2, were obtained from Dr. Ping Chen, Otogenetics Corporation (Qian et al., 2007), and crossed to *Scrib^rumz/+^* mice to produce *Scrib^rumz/+^; Vangl2^GFP/+^* heterozygotes. B6.129(Cg)-*Gt(ROSA)26Sortm4(ACTB-tdTomato,-EGFP)Luo*/J (*mT/mG*) mice carrying floxed membrane-targeted Tomato fluorescent protein followed by membrane-targeted eGFP (Muzumdar et al., 2007) were obtained from The Jackson Lab and maintained as homozygotes in our colony. All animals were genotyped with PCR using the primers listed in Supplementary Table 1 and the genotype of *Scrib^Crc^* and *Scrib^rumz^* embryos were confirmed by sequencing. Crosses were set-up between males and females of desired strains and the morning a plug was identified was designated as E0.5. Embryos were dissected at the desired stage in Whole Embryo Culture Medium (WECM) (Yen et al., 2009). All animal use protocols were in compliance with PHS and USDA guidelines for laboratory animal welfare and reviewed and approved by the University of Virginia Institutional Animal Care and Use Committee.

### Live imaging

To view CE and cell behavior, male *Scrib^rumz/+^; mT/mG* mice, were mated to female *Scrib^rumz/+^; Sox2Cre* mice in order to induce recombination and expression of membrane eGFP in all cells of all embryos, due to the maternal effect of the Sox2Cre (Hayashi et al., 2003). Live imaging was carried out as previously described (Williams et al., 2014). Briefly, embryos were dissected at the 0-3 somite stage and cultured in a mixture of 50% WECM and 50% rat serum. Embryos were mounted distal tip down in a glass-bottomed chamber, covered with culture medium which was then overlaid with mineral oil to prevent evaporation, and placed on the Zeiss 510 or Leica SP8 confocal microscope in the University of Virginia Keck Center (PI: Ammasi Periasamy; NIH- RR025616) in an environmental chamber at 37°C and 5% CO_2_ gas. For CE and cell behavior analysis, embryos were imaged for 6-10 hours at an interval of 6min at 40x with Z-stacks of 10-15 planes captured at 2μm intervals. All embryos were lysed and genotyped either by PCR or by sequencing after imaging.

### Image Analysis

In order to best visualize the Z-plane of interest in the tissue for CE analysis confocal stacks were aligned by concatenating single optical sections across time. Movies were also aligned in the X and Y axes using the Stackreg plugin in ImageJ to eliminate whole embryo drift from cell tracking results. Rectangular distortion diagram boxes, oriented with their long axis parallel to the AP axis of the neural plate, were generated from cell tracking of midline neural plate cells throughout live imaging movies. ImageJ was then used to calculate the CE index, which is represented by the change in the aspect ratio of the length of the AP axis (length of boxes) over the width of the ML axis (width of the boxes). The change in AP length represents the amount of elongation, while the change in ML width represents the amount of convergence. For intercalation analysis midline cells were manually tracked and observed in clusters of approximately 10 cells in order to evaluate intercalation frequency and the polarity of separation of neighboring cells. Individual cells were measured to analyze basal and apical area and apical constriction index (ACI), ratio of basal cell area over apical cell area, as well as basal orientation by fitting an ellipse to each cell at both the basal and apical surfaces

### Immunofluorescence staining

For immunofluorescence whole mount staining of E8.5 embryos were fixed in 3.7% paraformaldehyde for 2 hours. Following fixation, embryos were washed overnight in tsPBS (0.5% triton + 0.1% saponin in PBS), blocked for 1 hour at 37°C in 10% goat serum in tsPBS, and incubated in the following primary antibodies diluted 1:100 (except phalloidin, 1:300) in blocking solution overnight at 4°C: aPKC (sc-216, SCBT), ZO-1 (21773-1-AP, proteintech), E-cad (610181, BD), N-cad (610920, BD), Par3 (07-330, Millipore), Par6 (sc-166405, Santa Cruz), rhodamine phalloidin (R415,Invitrogen), Myosin IIA (PRB-440P,Covance), Myosin IIB (PRB-445P,Covance), and phospho-Myosin Light Chain (3674, Cell Signaling). Following primary incubation, embryos were washed in tsPBS and incubated in the following Alexa-conjugated secondary antibodies 1:200 in blocking solution overnight at 4°C: Alexa-Fluor donkey anti-rabbit IgG 488 (A21206, Life Tech), Alexa-Fluor goat anti-mouse IgG 546 (A11030, Life Tech), Alexa- Fluor goat anti-rabbit IgG 647 (A21245, Life Tech), and Alexa-Fluor donkey anti-mouse IgG 488 (A21202, Life Tech). Embryos were washed with tsPBS and imaged on Zeiss 780 confocal microscope in the University of Virginia Keck Center (PI: Ammasi Periasamy; NIH-ODO16446). Zen software was used to acquire images and ImageJ was used to analyze images.

### Western Blot

For analysis of whole cell lysates, whole embryos were lysed directly in Laemmli sample buffer, subjected to SDS-PAGE, transferred to nitrocellulose membrane, blocked in 1% FG in TBST 1 hour at 4°C, and analyzed by Western blotting with Scrib primary antibody (A01651, Boster Bio) 1:1000 in blocking solution incubated overnight at 4°C. Protein was visualized with the following IRDye conjugated secondary antibodies 1:20,000 in TBST + 0.01% SDS for 45min at room temperature: rabbit680rd and mouse800cw (LiCor). Tubulin was used as the loading control. LiCor Image Studio software was used to acquire and analyze images.

### Statistics

PAST software (https://www.nhm.uio.no/english/research/infrastructure/past/) was used to make rose diagrams and to perform the associated circular statistics including circular mean and Mardia-Watson-Wheeler Test to test for equal distributions. Graphpad Prism 8 software was used to create all other graphs and was used to perform all other statistical analyses including Fisher’s Exact test, t-test, one-way ANOVA and two-way ANOVA tests. Significance was determined by p<0.05 and “ns” is used to denote not significant (p>0.05).

## Results

### Scrib^rumz^ and Scrib^Crc^ mutants exhibit similar phenotypes of open neural tube and lack of axial turning

The *Scrib^rumz^* mutant has not been characterized in detail, so to compare effects of this mutation, which reduces Scrib expression and localization ((Dow et al., 2007) and Supplemental Figure 1), and the *Scrib^Crc^* truncation mutation, which is an effective null ((Murdoch et al., 2014) and Supplemental Figure 1), E11.5 embryos were dissected and imaged to examine phenotypes (Figure 1A). The overall incidence of NTD was the same between the two mutants, 92.3%, while only 6.9% of *Scrib^rumz/+^* and 6.1% of *Scrib^Crc/+^* heterozygous embryos exhibit NTD (Figure 1A, B). The most dominant NTD observed was CRN with 81% of *Scrib^rumz/rumz^* and 88% of *Scrib^Crc/Crc^* mutant embryos displaying CRN (Figure 1A, B). However, a finer-grained analysis of the phenotypes revealed differences between the two mutant strains. Phenotypes were further classified as CRN, exencephaly, turning defects, kinked tail, and developmental delay (Supplemental Table 2). While both *Scrib^rumz/rumz^* and *Scrib^Crc/Crc^* exhibit a significantly larger percentage of embryos with CRN and turning defects, *Scrib^rumz/rumz^* display more embryos with exencephaly while *Scrib^Crc/Crc^* display a significantly larger percentage of embryos with kinked tails compared to their respective wildtype littermates. Minor defects were observed in 3-25% of wildtype and heterozygous embryos likely due to the mixed backgrounds of the transgenic lines in our colony. Furthermore, *Scrib^Crc/Crc^* and *Scrib^rumz/rumz^* embryos both show incomplete penetrance of NTD, (Figure 1 and (Murdoch et al., 2014; Murdoch et al., 2001b; Zarbalis et al., 2004)), likely due to the mixed genetic background of our transgenic lines which has been previously demonstrated to influence the severity of NTD in *Scrib^Crc/Crc^* mutants (Murdoch et al., 2014).

**Figure 1.**
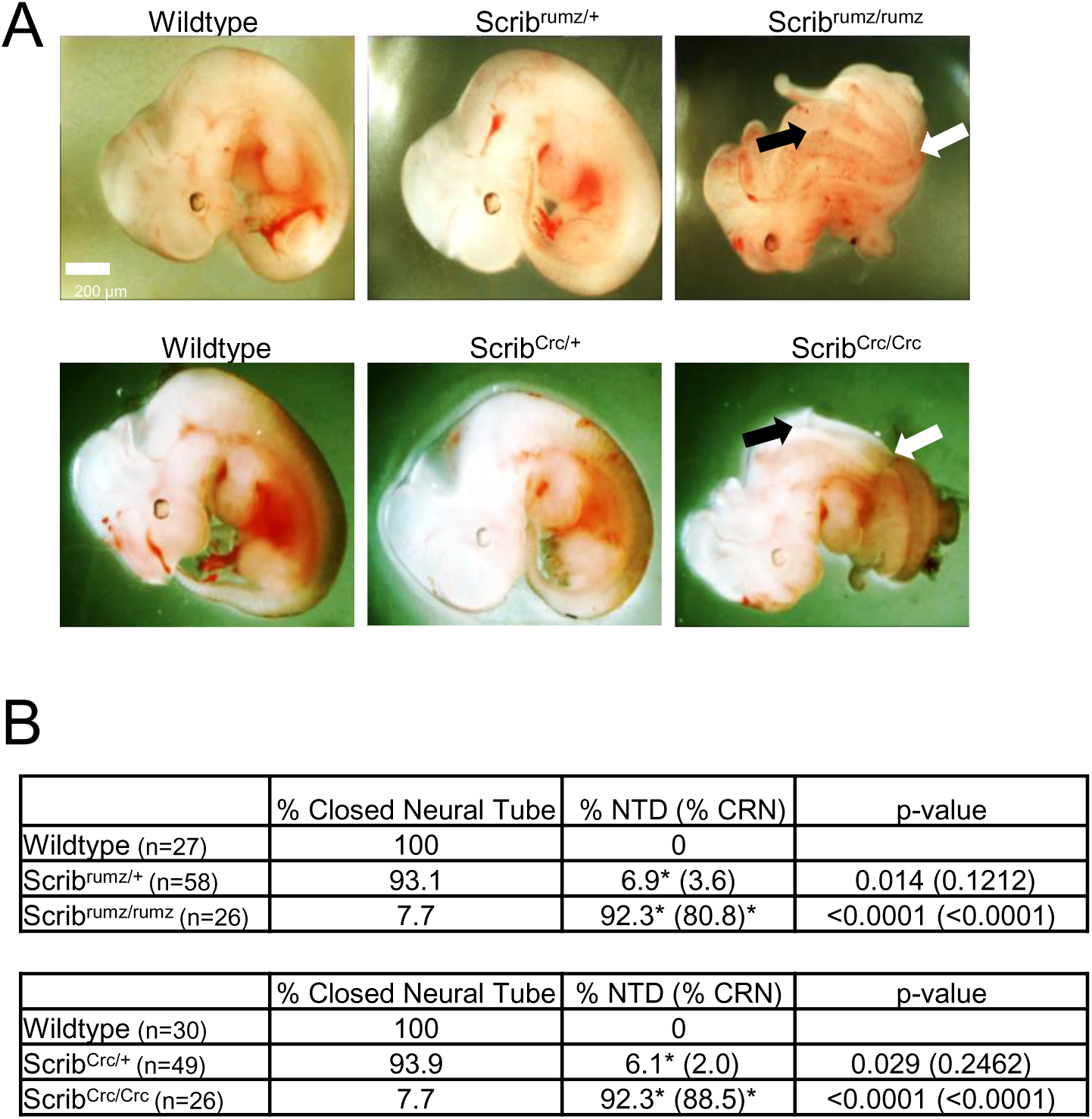
*Scrib* mutants display open neural tube and turning defects. A) Bright field images of E11.5 embryos, scale bar = 200µm, black arrows indicate CRN and white arrows indicate turning defect. B) Table displays percent of embryos with either closed neural tube or neural tube defects (NTD). NTD include CRN and exencephaly phenotypes with the percentage of embryos with CRN, the dominant NTD, displayed in parentheses. Significance is marked with * and was determined by Fisher’s Exact test compared to wildtype littermates. p-values for NTD are displayed in the last column of the table with the p-value for CRN alone in parentheses. n represents the number of embryos analyzed.

### Scrib mutants exhibit different genetic interactions with Vangl2^Lp^ and Vangl2^KO^

Both *Scrib^rumz^* and *Scrib^Crc^* mutations interact genetically with the *Vangl2^Lp^* mutation to cause CRN (Murdoch et al., 2001b; Zarbalis et al., 2004). To more completely characterize the phenotypes resulting from *Scrib* and *Vangl2* genetic interactions, phenotypes of *Scrib^rumz^; Vangl2^Lp^* and *Scrib^Crc^; Vangl2^Lp^* embryos were compared and the genetic interactions between *Scrib* mutants and Vangl1 knockout (*Vangl1^KO^*) and Vangl2 knockout (*Vangl2^KO^*) were examined at stage E11.5 (Figure 2A). Consistent with previous studies *Scrib^rumz/+^; Vangl2^Lp/+^* and *Scrib^Crc/+^; Vangl2^Lp/+^* double heterozygous embryos display a small but significant increase in the percentage of embryos with NTD, 27% and 16% respectively, compared to the respective single heterozygotes (Figure 2B, (Murdoch et al., 2001b)). Interestingly, loss of one allele of either Vangl1 (*Vangl1^KO/+^*) or Vangl2 (*Vangl2^KO/+^*) in addition to mutation of one Scrib allele has no significant effect on NTD as no *Scrib^rumz/+^*; *Vangl1^KO/+^* embryos exhibit NTD and only 4.8% of *Scrib^rumz/+^*; *Vangl2^KO/+^* embryos display NTD. However, *Scrib^rumz/+^*; *Vangl1^KO/+^; Vangl2^KO/+^* triple heterozygotes do display a significant increase in NTDs with approximately 13% exhibiting NTD, although this is still smaller than the percentage of *Scrib^rumz/+^; Vangl2^Lp/+^* embryos (Figure 2B). Similar results were observed in *Scrib^Crc/+^*; *Vangl1^KO/+^*, *Scrib^Crc/+^*; *Vangl2^KO/+^*, and *Scrib^Crc/+^*; *Vangl1^KO/+^; Vangl2^KO/+^* embryos (data not shown).

**Figure 2.**
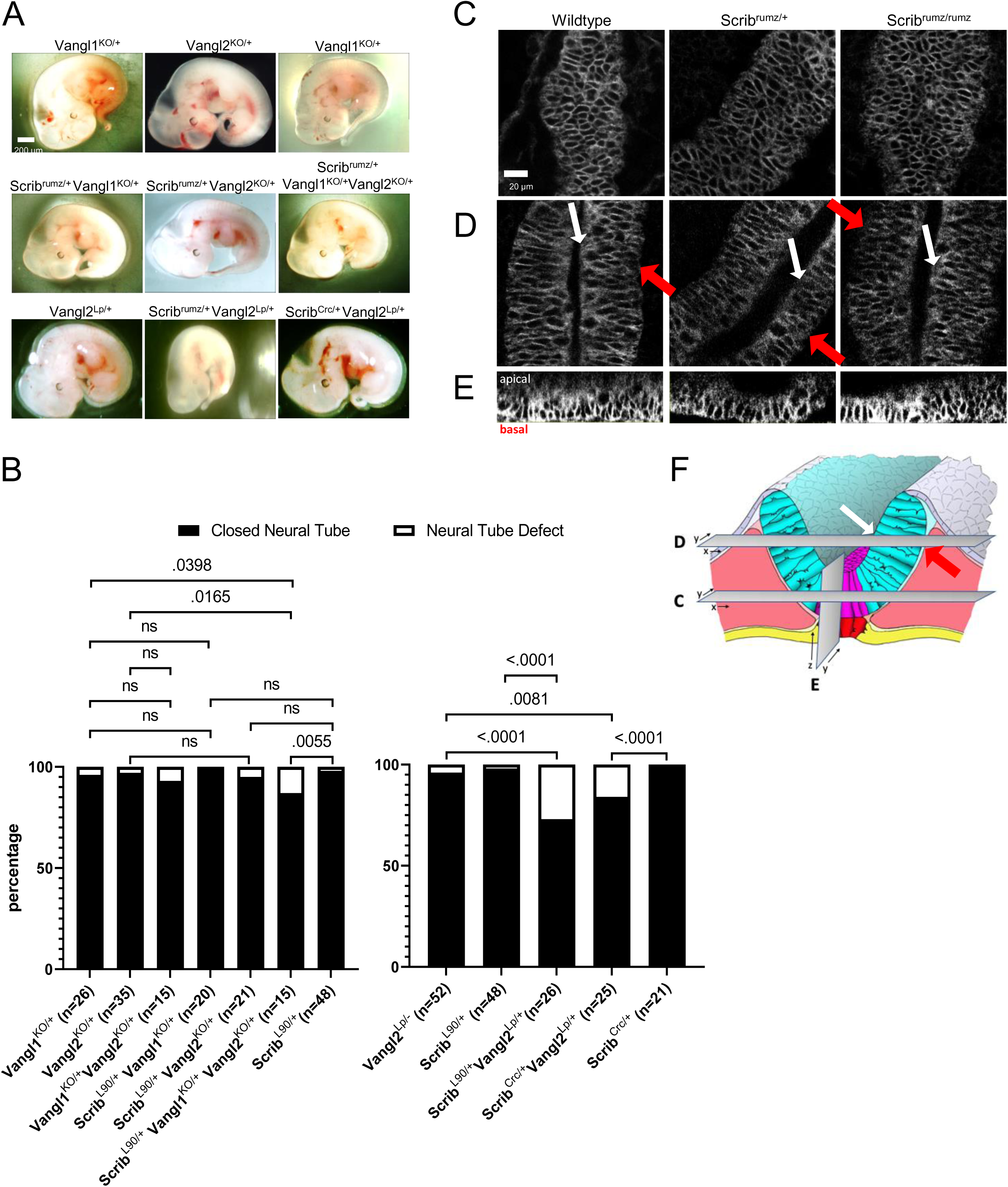
*Scrib* interacts genetically with *Vangl2^Lp^*. A) Bright field images of E11.5 embryos, scale bar = 200µm. B) Graph displaying the percentage of embryos with closed neural tube or neural tube defects in E11.5 embryos. p-values were determined by Fishers Exact test (significance p <0.05) and ns denoting not significant (p >0.05). n represents the number of embryos analyzed. C-F) Confocal images of Vangl2 localization in *Scrib^rumz^* E8.5 embryos: in the floorplate (C), neural folds where white arrows indicate the apical surface and red arrows indicate the basal surface (D), and orthogonal ZY projections of the floorplate oriented with apical surface on top and basal surface on bottom (E). scale bar = 20µm. F) Schematic indicating the relative position of views in C, D, and E in the neural tissue.

To observe Vangl2 localization when *Scrib* is mutated we utilized the Vangl2^GFP/GFP^ transgenic mouse line carrying a BAC expressing GFP tagged Vangl2 (Qian et al., 2007). Embryos from *Scrib^rumz/+^; Vangl2^GFP/+^* double heterozygote crosses were dissected and fixed at E8.5 and the neural tissue was imaged (Figure 2F). Vangl2 was similarly localized to the cell membrane throughout the tissue in wildtype, *Scrib^rumz/+^* and Scrib^rumz/rumz^ embryos specifically in both the floorplate (Figure 2C, E) and in the wings of the neural folds (Figure 2D), suggesting that mutation of *Scrib* does not affect the membrane localization of Vangl2 protein. These results in *Scrib^rumz^* tissue are different from previously published data in *Scrib^Crc^* mice lacking Scrib expression where Vangl2 localization is increased at the apical membrane in *Scrib^Crc/Crc^* neural tissue (Kharfallah et al., 2017).

Overall, these data are consistent with previous studies showing that *Scrib* genetically interacts with *Vangl2^Lp^* (Murdoch et al., 2001b; Zarbalis et al., 2004), however they highlight that the genetic interaction with *Vangl2^Lp^* is not solely due to the combined decrease in normal Scrib and Vangl2 proteins but must be due to additional effects on other protein(s) by the *Vangl2^Lp^* mutation. Not only are the phenotypes that result from genetic interactions of *Scrib^rumz/+^* and *Vangl2^KO/+^* less severe, but they are qualitatively different from those seen in *Scrib^rumz^; Vangl2^Lp^* embryos (data not shown), and the additional loss of one allele of *Vangl1* in *Scrib^rumz/+^*; *Vangl1^KO/+^; Vangl2^KO/+^*embryos does not phenocopy the genetic interaction with *Vangl2^Lp^*.

### Scrib affects convergent extension of the neural tube

To determine whether the NTD observed in *Scrib* mutants could be attributed to inhibition of convergent extension (CE), we crossed *Scrib^rumz^* mice with the *mT/mG* line, and used fluorescently labeled E8.0 (1-3 somites) embryos in live confocal imaging for 6-10 hours to visualize tissue shape changes and overall elongation during NTC. Distortion diagrams drawn from relative cell positions tracked over time were used to quantify CE of the neural plate. In wildtype and *Scrib^rumz/+^* heterozygote movies the tissue elongates and narrows over time, while *Scrib^rumz/rumz^* mutant tissue underwent less CE (Figure 3A). The overall change in CE index (aspect ratio at the end of the movie over the aspect ratio at the start) per hour in wildtype and *Scrib^rumz/+^* embryo movies is approximately 2% and 1.5% and is significantly decreased to -0.5% in *Scrib^rumz/rumz^* embryo movies (Figure 3B).

**Figure 3.**
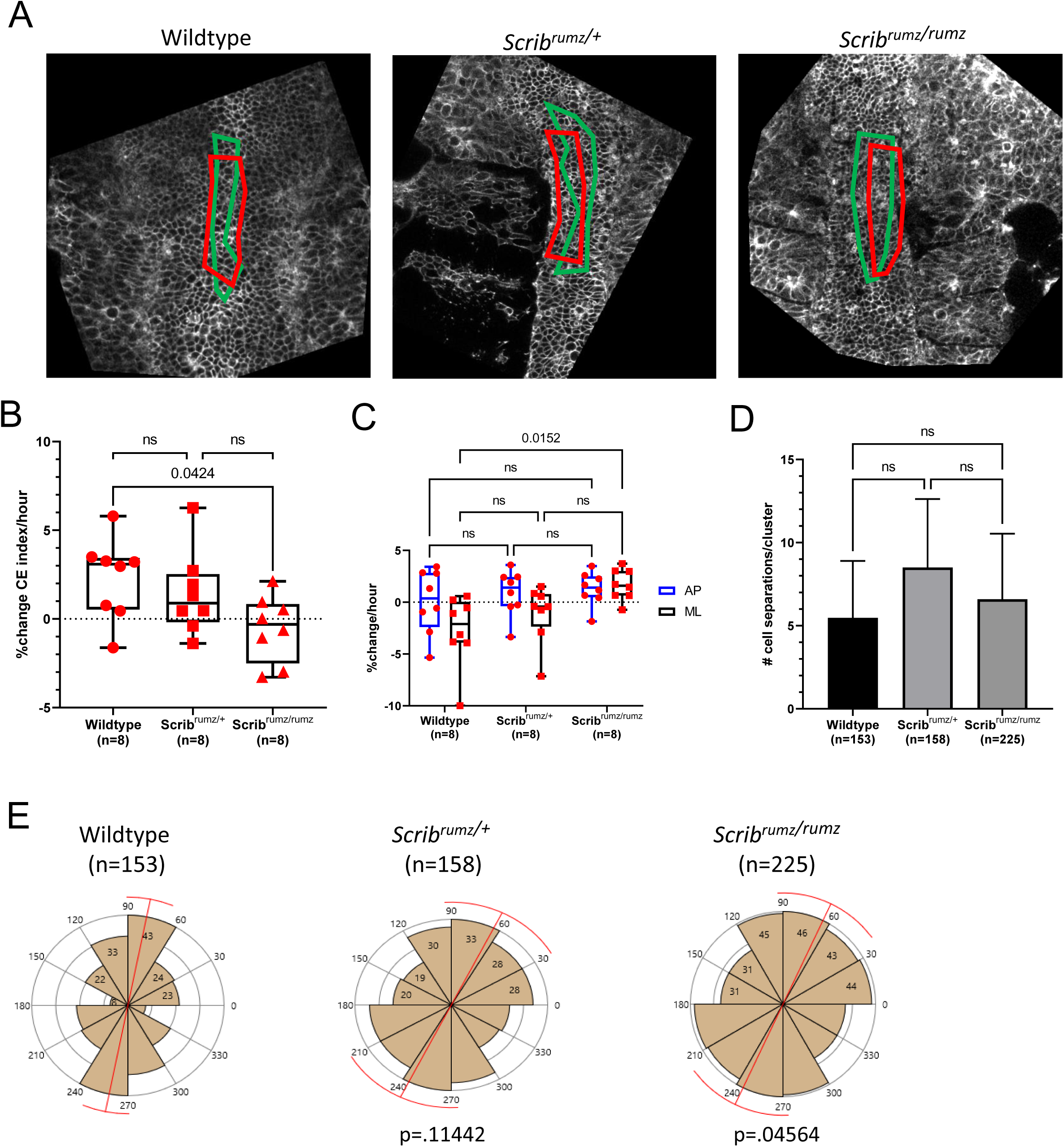
*Scrib^rumz^* mutation affects convergent extension and polarity of intercalation. A) Representative images from live imaging movies showing distortion diagrams drawn at the start (red boxes) and end (green boxes) of movies. B) Box and whisker plot of percent change in CE index (aspect ratio at the end of the movie over the aspect ratio at the start) per hour over 6-10 hours of live imaging movies of E8.0 mouse embryos fluorescently labeled with *mG/mT*. Error bars represent min and max and significance was determined by one-way ANOVA (p<0.05) with ns denoting not significant (p>0.05). n represents the number of embryos analyzed. C) Graph summarizing percent change per hour in AP and ML dimensions of live imaging movies. Error bars represent min and max and significance was determined by two-way ANOVA (p<0.05) with ns denoting not significant (p>0.05). n represents the number of embryos analyzed. D) Graph summarizing total number of cell separations observed in the neural tissue. Error bars represent standard deviation and significance was determined by one- way ANOVA (p<0.05) with ns denoting not significant (p>0.05). n represents the number of intercalations analyzed. E) Rose plots displaying the distribution of AP (60-120°), diagonal (30-60° and 120-150°), and ML (0-30° and 150-180°) neural cell separations. Red bar represents circular mean and p values were determined by Mardia-Watson- Wheeler Test for equal distributions comparing *Scrib^rumz/+^* or *Scrib^rumz/rumz^* to wildtype (significance p<0.05). n represents number of intercalations analyzed and 8, 6, and 9 embryos were analyzed for wildtype, *Scrib^rumz/+^* and *Scrib^rumz/rumz^* respectively.

When broken down to axial components, we found no significant difference in the amount of elongation (AP) between the genotypes, but observed a significant difference in the amount of convergence (ML) (Figure 3C). The wildtype neural plate narrows (- 2.7% change per hour) while the *Scrib^rumz/rumz^* neural plate widens (+1.5% change per hour). The *Scrib^rumz/+^* neural plate does narrow although to a lesser degree (-1.0% change per hour) (Figure 3C). The number of cells within the distortion diagram area remain the same at the start and end of the movies in wildtype, *Scrib^rumz/+^*, and *Scrib^rumz/rumz^* (Supplemental Figure 2A), while the overall area of the designated distortion diagram decreases as the movies progress in wildtype but increases in *Scrib^rumz/+^* and *Scrib^rumz/rumz^* movies (Supplemental Figure 2B). These results suggest that changes in tissue shape and lack of CE are not due to changes in cell number but instead due to differences in cell intercalation and cell shape.

### Scrib regulates mechanisms of cell rearrangement and polarity of intercalation

CE results from cells intercalating mediolaterally to extend the tissue in the AP direction (Keller, 2002; Keller and Sutherland, 2020; Sutherland et al., 2020), and we have shown previously that defects in the polarity or frequency of cell intercalation lead to decreased CE (Williams et al., 2014). The frequency and polarity of cell intercalation in embryos from *Scrib^rumz/+^* heterozygote crosses were assessed by live imaging. Cells in the midline were tracked throughout the movies of *mT/mG* fluorescently labeled E8.0 (1-3 somites) described above, and for each embryo, rearrangements were analyzed in several clusters of approximately 10 cells each. No significant differences in the total number of cell separations per cluster were observed between wildtype, *Scrib^rumz/+^* and *Scrib^rumz/rumz^* embryos (Figure 3D). However, mutation of *Scrib* had a strong effect on the polarity of cell intercalation. To determine the polarity of cell intercalations the angle of separation of neighboring cells was measured relative to the AP axis of the embryo, with the assumption that separation of neighboring cells along the AP axis results from a mediolateral intercalation, while a separation along the ML axis represents an AP intercalation. The angles were then plotted to show the orientation of cell separations and a Mardia-Watson-Wheeler Test was used to test for equal distributions. The majority of wildtype cell separations occurred in the AP direction, indicating mediolaterally polarized intercalation. Similarly, cells in the *Scrib^rumz/+^* tissue primarily separate in the AP direction determined by p>0.05 compared to wildtype, whereas *Scrib^rumz/rumz^* cells separations were significantly different from wildtype (p<0.05) (Figure 3E).

The rearrangements were classified as resulting from rosette resolution, T1 process, single cell intercalation, or from cell division (Figure 4A and (Williams et al., 2014)). The relative frequency of the various types of cell rearrangements was similar between wildtype, *Scrib^rumz/+^* and *Scrib^rumz/rumz^* embryos (Figure 4B). However, we observed distinct effects of the *Scrib^rumz^* mutation on the process of rosette resolution, with many rosettes forming and then resolving back in the original direction, and others forming, resolving, and then reforming. To characterize rosette formation and resolution in more depth, all rosettes observed were classified as either not resolved (a rosette formed and remained in the rosette configuration until the end of the movie), resolved by junctional rearrangement (a rosette forms and coordinated junctional rearrangement leads to neighbor exchange), or resolved by division (a rosette forms and division of cells within or near the rosette disrupts its configuration) (Figure 4C). We further classified the rosettes that resolved by junctional rearrangement into two groups: those in which the cells go from orientation along one axis to rosette formation and then resolve into orientation along the other axis (productive junctional rearrangement), and those in which the cells go from orientation along one axis to rosette formation and then resolve back along the original axis (unproductive junctional rearrangement) (Figure 4C). A final group, only observed in *Scrib^rumz/+^ and Scrib^rumz/rumz^* embryos, were rosettes that repetitively formed and resolved unproductively (pulsing rosettes) (Figure 4C). These categories differed significantly between the three genotypes (Figure 4D): the frequency of productive junctional rearrangement went down in the *Scrib^rumz/+^* embryos, and was gone in the mutant, while the frequency of unproductive junctional rearrangements and pulsing rosettes went up in the *Scrib^rumz/+^* embryo, and increased further in the mutant.

**Figure 4.**
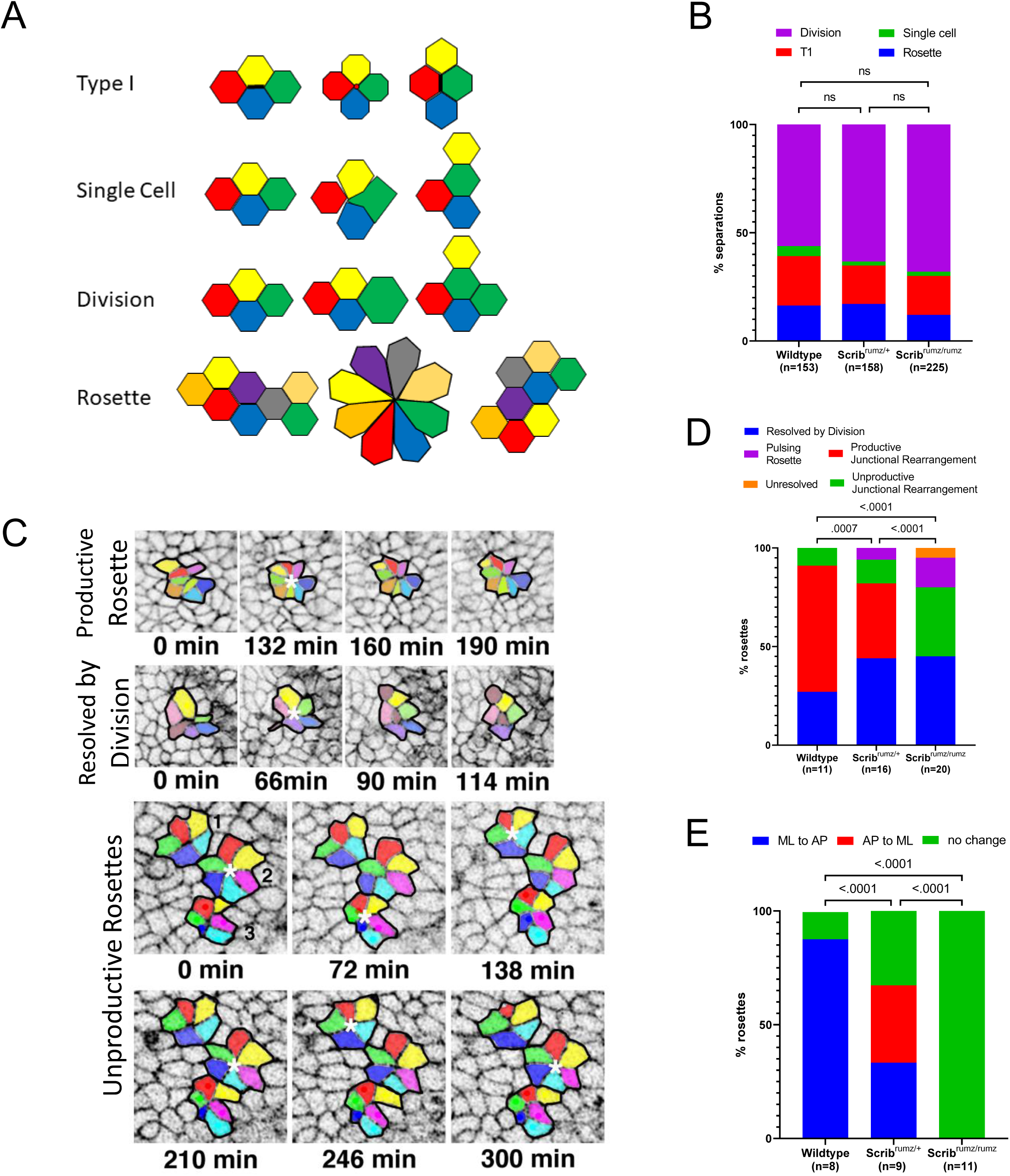
*Scrib^rumz^* mutation affects rosette resolution. A) Schematic summary of each type of cell separation mechanism. B) Defects in cell rearrangement mechanisms in neural tissue were assessed by analyzing the percentage of cell intercalations arising from cell division, single cell intercalation, T1 process, and rosette resolution. ns denotes not significant (p>0.05) and was determined by Fisher’s Exact test. n represents the number of intercalations analyzed. C) Representative images of types of rosette resolution. In the panels representing unproductive rosettes, rosette 1 and 2 represent pulsing rosettes, and rosette number 3 shows an unproductive junctional rearrangement resolution. Rosettes are marked with white * and each rosette cluster is outlined in black. D) Graph represents the mechanisms of rosette resolution. p values were determined by Fisher’s Exact test (significance p<0.05) and n represents the number of rosettes analyzed. E) Graph representing the changes in cell cluster orientation during rosette resolution. p values were determined by Fisher’s Exact test (significance p<0.05) and n represents the number of rosettes analyzed that resolved by a mechanism other than division (pulsing, productive junctional rearrangement, unresolved, and unproductive junctional rearrangement).

In addition, the specific polarity of formation and productive resolution of rosettes was strongly affected by the *Scrib^rumz^* mutation. In wildtype embryos the majority of rosettes formed along the ML axis and resolved along the AP axis, whereas in *Scrib^rumz/+^* embryos there was an equal frequency of formation along the ML and AP axes and resolution along the other axis (Figure 4E). Furthermore, no *Scrib^rumz/rumz^* rosettes resolved productively by junctional rearrangement with about 55% of rosette resolutions resulting in unproductive separations and the remaining resolutions taking place by division (Figure 4E). Importantly, none of the rosettes formed in *Scrib^rumz/rumz^* tissue led to a change in the orientation of the cell clusters, emphasizing the lack of productivity of rosette intercalations in *Scrib^rumz/rumz^* embryos (Figure 4E). Together these data underline an important role for Scrib in determining the polarity of intercalation as well as in promoting rosette resolution and suggest that defects in polarized cell intercalation contribute to impaired tissue shape changes and CE in *Scrib^rumz/rumz^* embryos.

### Scrib promotes apical constriction

Cells must transform from columnar to wedge shape in order for the neural tube to bend (Inoue et al., 2016; McShane et al., 2015; Smith et al., 1994). Our previous data show that floorplate cells normally increase area and elongate basally along the mediolateral axis, while decreasing area (constricting) and remaining more rounded apically (Williams et al., 2014). We observed normal basal elongation and orientation of cells in wildtype and *Scrib^rumz/+^* embryos at the midline, however, the basal ends of neural cells at the midline of *Scrib^rumz/rumz^* embryos did not elongate, and their long axis was randomly oriented relative to the AP axis of the embryo (Figure 5A). Rose plots representing the orientation of the long axis of the basal ends of cells show predominant orientation along the ML axis (0-180°) in both wildtype and *Scrib^rumz/+^* embryos, whereas *Scrib^rumz/rumz^* cells are not oriented in a particular direction as shown by the uniform rose plot (Figure 5A). Cell wedging occurred in wildtype embryos at the midline as apical cell area decreased (Figure 5B), while the apical constriction index (ACI), the ratio of basal cell area over apical cell area, increased (Figure 5C). Surprisingly, in *Scrib^rumz/+^* midline cells apical cell area increased and the ACI decreased suggesting that the apical end is not constricting but instead getting larger, and the cells are becoming more columnar (Figure 5B, C). Cell area in *Scrib^rumz/rumz^* neural cells at the midline does not change significantly at the apical end resulting in a decrease in ACI (Figure 5B, C). Thus, *Scrib* mutants do not transform from columnar to wedge shaped cells as required for neural plate bending, suggesting a role for Scrib in regulating apical constriction during NTC. Furthermore, the decreased apical constriction in heterozygous embryos suggests that the mutation may have some dominant effects on this process and may contribute to the minor NTDs we observe in *Scrib^rumz/+^* embryos.

**Figure 5.**
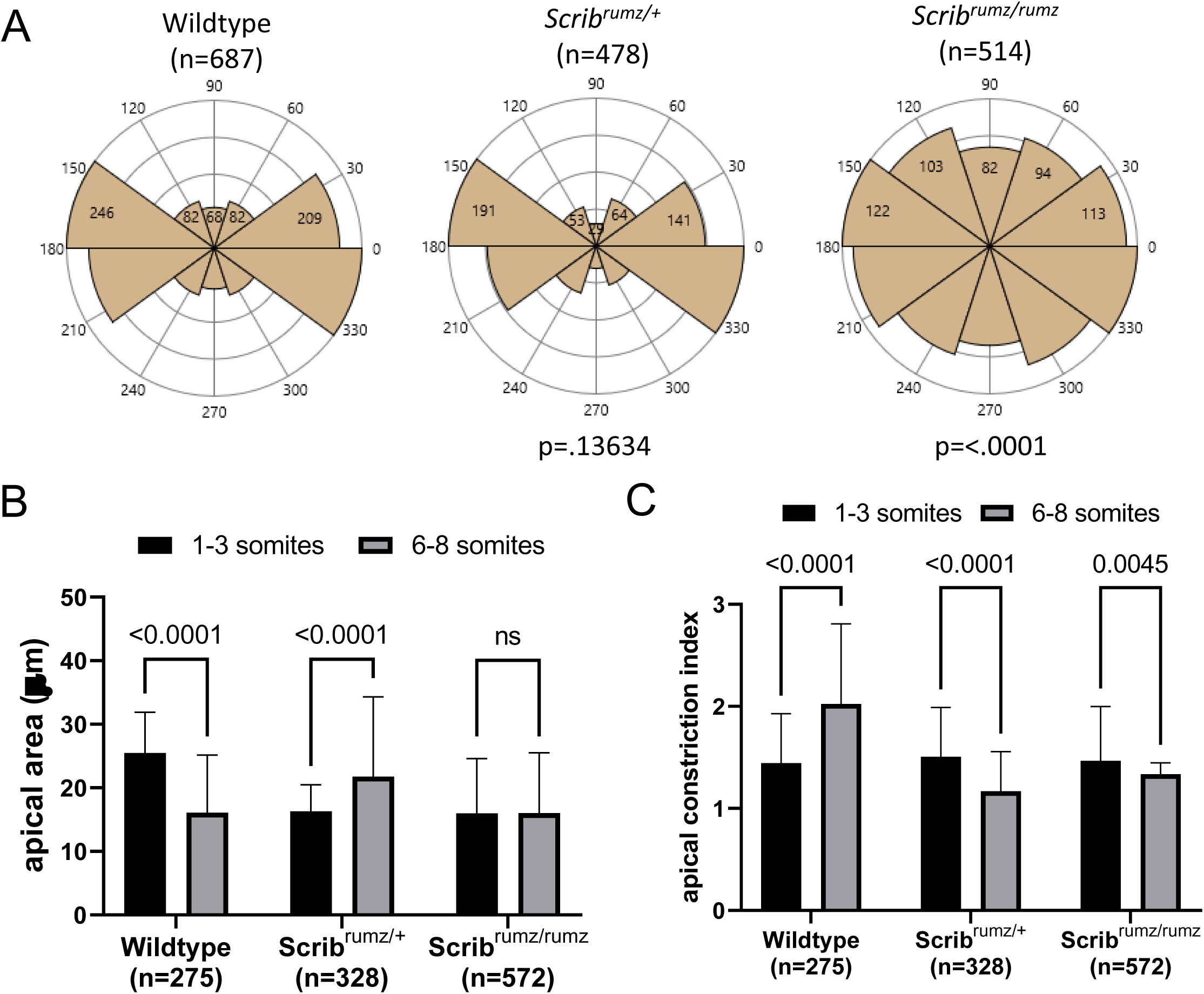
Apical constriction is disrupted in *Scrib^rumz^* mutants. A) Rose diagrams represent basal cell orientation where 90-270° indicates the AP axis and 0-180° indicates the ML axis. p-values were determined by Mardia-Watson-Wheeler Test for equal distributions comparing *Scrib^rumz/+^* or *Scrib^rumz/rumz^* to wildtype (significance p<0.05). n represents number of cells analyzed and 8, 3, and 5 embryos were analyzed for wildtype, *Scrib^rumz/+^* and *Scrib^rumz/rumz^* respectively. B) Graph indicates apical cell area at the start (1-3 somites) and end (6-8 somites, 6-10 hours) of live imaging movies of E8.0 embryos. Error bars represent standard deviation. p-values were determined by two-way ANOVA (significance p<0.05) and ns denotes not significant (p>0.05). n represents number of cells analyzed. C) Graph displays apical constriction index (ACI), the ratio of basal cell area over apical cell area, at the start and end of live imaging movies. Error bars represent standard deviation. p-values were determined by two-way ANOVA (significance p<0.05). n represents number of cells analyzed.

### Localization and expression of junctional proteins is affected by Scrib mutation

Cell rearrangements and cell shape changes rely on cell-cell interactions, shortening of junctions, and formation of new junctions (Rauzi et al., 2010; Shindo, 2018). Given Scrib’s role in maintaining junctional composition, mediating junctional remodeling, and regulating E-cadherin retention and recycling at the adherens junction (Bonello and Peifer, 2018; Lohia et al., 2012), it is possible that Scribble regulates CE and apical constriction by influencing the localization and expression of junctional proteins. One component of the Atypical Protein Kinase C (aPKC) complex, Partitioning defective 6 (Par6) which is normally enriched at the apical surface and at tight junctions is decreased at the apical surface in *Scrib^rumz/rumz^* cells but unchanged in *Scrib^rumz/+^* cells (Figure 6A, B). Localization of partitioning defective 3 (Par3), another component of the aPKC apical polarity complex, is decreased in both *Scrib^rumz/+^* and *Scrib^rumz/rumz^* tissue compared to the wildtype neural plate (Figure 6A, B). aPKC expression remains unchanged between the three genotypes (Figure 6A, B). The tight junction protein Zonula Occludin-1 (ZO-1) is decreased at junctions in both *Scrib^rumz/+^* and *Scrib^rumz/rumz^* cells as compared to wildtype (Figure 6A, B). Expression of both N-cadherin (N-cad) and Ecadherin (E-cad) is unchanged in *Scrib^rumz/+^* tissue, but decreased in *Scrib^rumz/rumz^* tissue as compared to wildtype (Figure 6A, B). Thus, *Scrib^rumz^* affects the expression and localization of several apical and tight junction proteins including Par6, Par3, and ZO-1 and disrupts retention of cadherin at the membrane confirming a role for Scrib in targeting proteins critical for cell interactions and junctional remodeling to the cell membrane during NTC.

**Figure 6.**
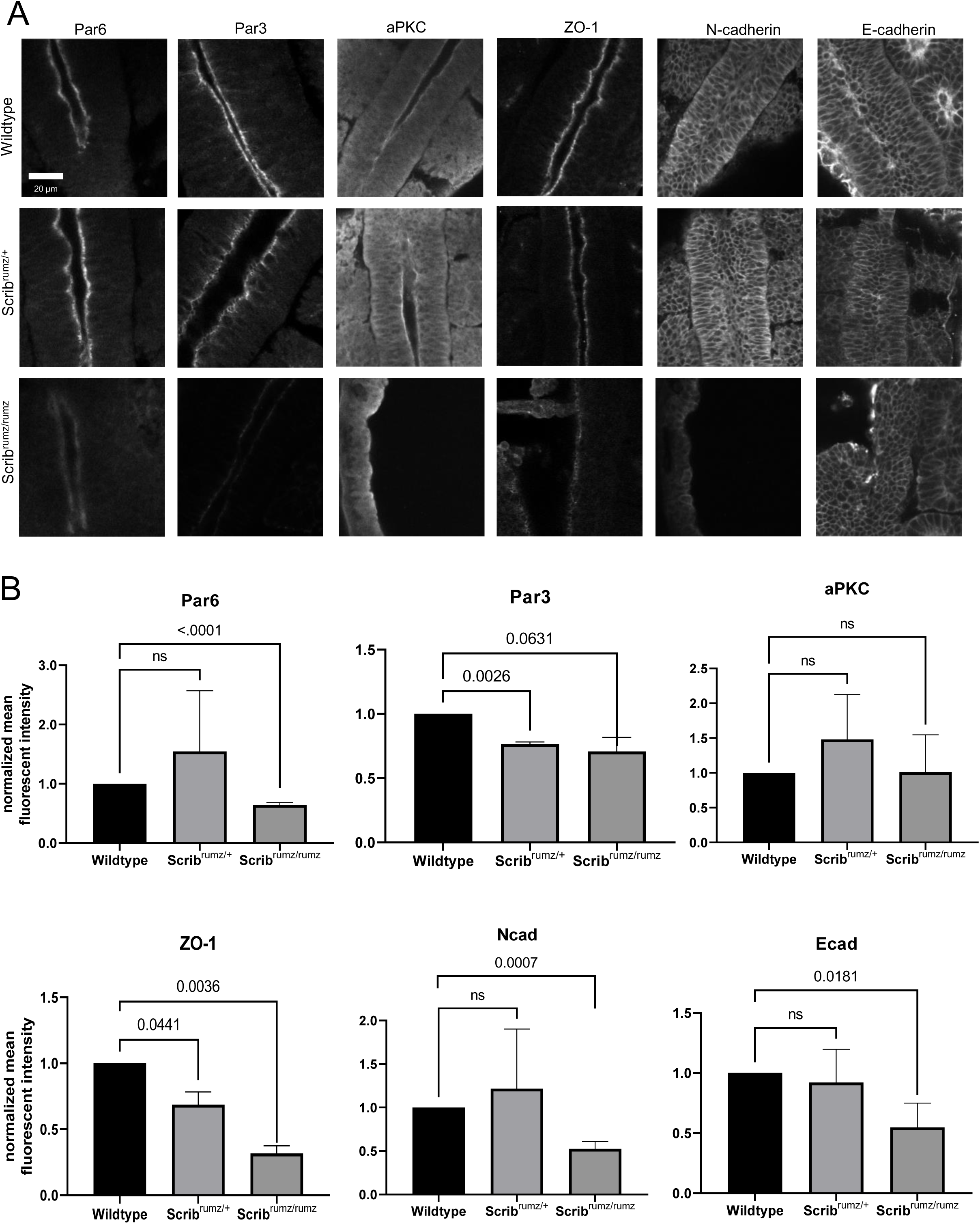
*Scrib^rumz^* mutants exhibit mislocalization of junctional proteins. A) Z slice of confocal images of whole mount staining for junctional proteins in E8.5 embryos. B) Line scans were used in ImageJ to measure mean fluorescent intensity at the apical junctions (aPKC, Par6, Par3, and ZO-1) or membrane (N-cad and Ecad). Normalized values are plotted with error bars represent standard deviation. p-values were determined by t-test compared to wildtype (significance p<0.05) and ns denotes not significant (p>0.05).

### Scrib regulates cytoskeletal proteins during NTC

Junctional shrinkage and rearrangement depends on actin and myosin dynamics, organization, and anchorage to adherens junctions ((Bertet et al., 2004; Blankenship et al., 2006; Fernandez-Gonzalez and Zallen, 2011; Rauzi et al., 2010) and reviewed in (Heer and Martin, 2017; Sutherland and Lesko, 2020)). Scrib is known to interact with several cytoskeleton-associated proteins such as βPIX, vimentin, and Lgl (Osmani et al., 2006; Phua et al., 2009; Raman et al., 2018), thus Scrib may play a role in regulating the linkage between the actomyosin network and the adherens junction in neural epithelial cells. Actin staining is very crisp at cell membranes in wildtype neural tissue and enriched at the apical surface; however, in *Scrib^rumz/+^* tissue actin localization at cell junctions is decreased significantly and is further decreased in *Scrib^rumz/rumz^* tissue compared to wildtype cells (Figure 7A, B). Similarly, non-muscle myosin IIB (MIIB) is decreased in *Scrib^rumz/rumz^* tissue at cell junctions compared to wildtype (Figure 7A, B). Surprisingly, non-muscle myosin IIA (MIIA) junctional expression is increased in *Scrib^rumz/rumz^* tissue compared to wildtype (Figure 7A, B). The overall activity of apical myosin is significantly decreased in both *Scrib^rumz/+^* and *Scrib^rumz/rumz^* cells as shown by loss of staining for phospho-myosin light chain (pMLC) (Figure 7A, B). Disruption of actin expression and both myosin expression and activity in *Scrib^rumz/rumz^* mutants suggest that changes in actomyosin dynamics may underlie the observed defects in junctional rearrangement and cell shape changes during NTC.

**Figure 7.**
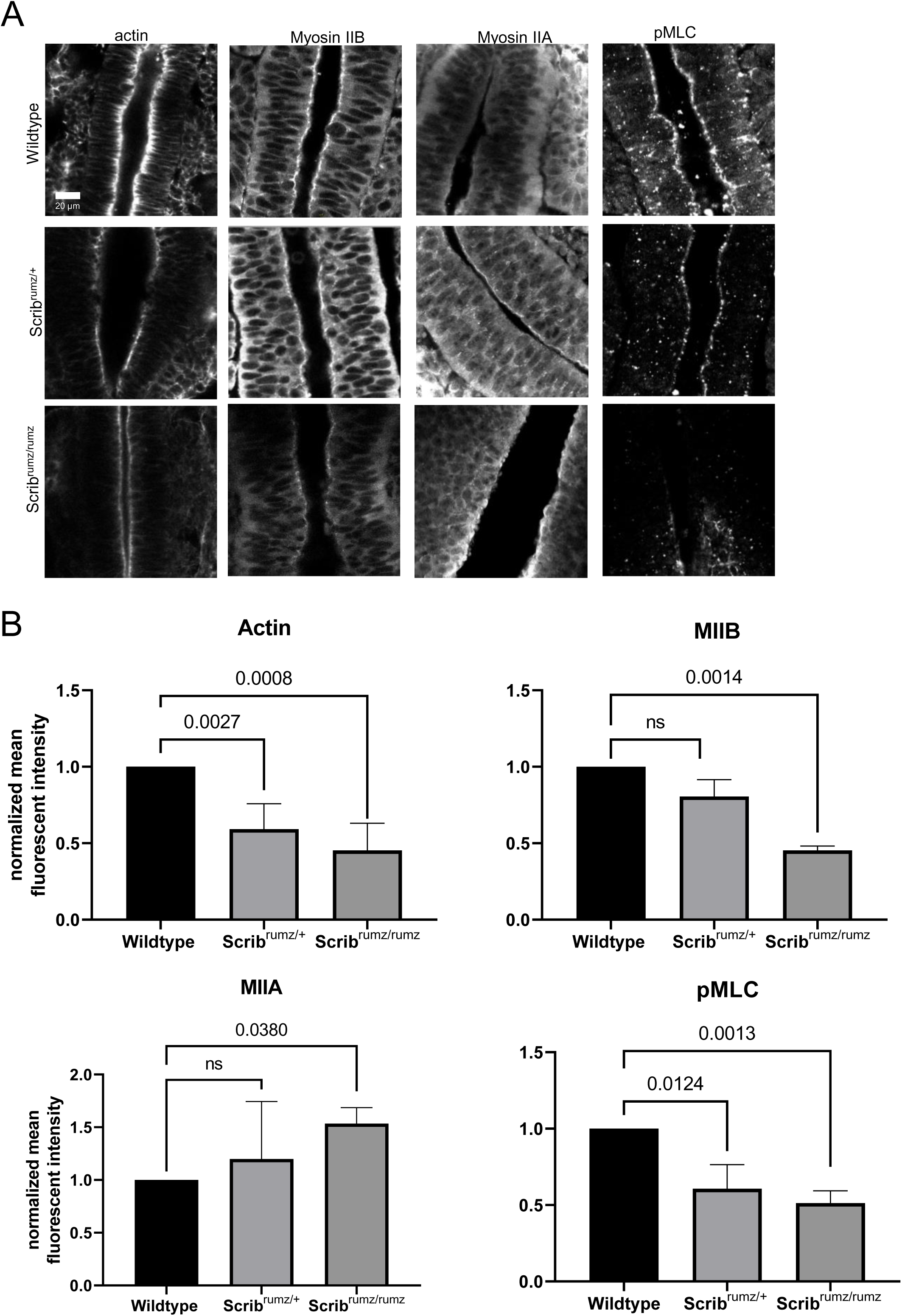
*Scrib^rumz^* mutants exhibit mislocalization of cytoskeletal proteins. A) Z slice of confocal images of whole mount staining for cytoskeletal proteins in E8.5 embryos. B) Line scans were used in ImageJ to measure mean fluorescent intensity at the apical junctions of neural cells. Normalized values are plotted with error bars represent standard deviation. p-values were determined by t-test compared to wildtype (significance p<0.05) and ns denotes not significant (p>0.05).

## Discussion

NTD occur in 1:500 pregnancies (Blencowe et al., 2018) resulting in birth defects such as anencephaly, spina bifida, and CRN, depending on where the failure of NTC occurs (Greene and Copp, 2014). Morphogenesis of the epithelial neural plate into the neural tube includes elongation and narrowing of the tissue through CE, and bending through apical constriction and cell wedging (Inoue et al., 2016; Keller et al., 2000; Keller and Sutherland, 2020; McShane et al., 2015; Sutherland et al., 2020). The PCP pathway has been shown to play a major role in orchestrating these events, as mutations in genes encoding PCP components lead to NTD (Greene and Copp, 2014; Wu et al., 2011; Zohn, 2020). Other proteins that affect PCP signaling similarly affect neural morphogenesis, and in particular, mutations in the gene encoding the protein Scrib cause NTD and interact genetically with mutations in *Vangl2*. However, the molecular and cellular mechanisms by which Scrib promotes NTC remain largely unknown. Here we show that Scrib is essential for both polarized cell intercalation during CE and cell shape changes that cause bending in the neural plate, and that it affects these processes in specific ways. Furthermore, we show that the Scrib mutation affects the composition of both tight and adherens junctions and the organization of the apical cytoskeleton. These studies clarify the role of Scribble in NTC, suggest future paths of investigation, and broaden our understanding of mammalian neural tube development.

### Scrib and PCP pathway interactions

Previous studies have shown that mutations in *Scrib* interact genetically with *Vangl2^Lp^* to cause CRN (Murdoch et al., 2014; Murdoch et al., 2001b; Zarbalis et al., 2004), which has been attributed to the direct interaction known to exist between Scrib and Vangl2 through the Scrib C-terminal PDZ domains (Belotti et al., 2013; Courbard et al., 2009; Kallay et al., 2006; Montcouquiol et al., 2006). Our observations of NTD in *Scrib^rumz/+^; Vangl2^Lp/+^* and *Scrib^Crc/+^; Vangl2^Lp/+^* are consistent with those previous results showing genetic interaction and identifying variable phenotypes in *Scrib^Crc/+^; Vangl2^Lp/+^* embryos (Murdoch et al., 2014; Murdoch et al., 2001b; Zarbalis et al., 2004). However, by examining double heterozygotes of either *Scrib^rumz/+^* or *Scrib^Crc/+^* and *Vangl2^KO/+^* as well as Vangl2 localization in *Scrib^rumz^* tissue, we show here that the genetic interaction between *Scrib* mutants and *Vangl^Lp^* is not simply due to heterozygosity for Scrib and Vangl2, as *Scrib^rumz/+^; Vangl2^KO/+^* embryos have less severe phenotypes than *Scrib^rumz/+^; Vangl2^Lp/+^* embryos. In principle, this could be attributed to compensation by Vangl1, given that Vangl1 and Vangl2 are very similar in structure and function (Gravel et al., 2010; Iliescu et al., 2011; Murdoch et al., 2001a; Torban et al., 2004) and that *Vangl2^Lp^* has a dominant negative effect on *Vangl1* (Yin et al., 2012). However, our results do not support this interpretation. Specifically, *Scrib^rumz/+^; Vangl1^KO/+^* embryos do not exhibit NTD, and a similar percentage of *Scrib^rumz/+^; Vangl2^KO/+^* double heterozygotes and *Scrib^rumz/+^; Vangl1^KO/+^*; *Vangl2^KO/+^* triple heterozygotes exhibit NTD. These observations suggest that the dominant effect of the *Vangl2^Lp^* mutation may be affecting a different, unknown, component of the PCP signaling pathway, or alternatively, that *Vangl2^Lp^* has a dominant negative effect on Scrib itself. The *Vangl2^Lp^* mutation inhibits trafficking of Vangl2 to the cell membrane (Merte et al., 2010), and it is possible that through direct or indirect interactions of Scrib and Vangl2, Scrib is trapped in the cytoplasm by Vangl2^Lp^ and therefore unable to complete necessary functions at the membrane for NTC.

### Scrib affects polarized cell intercalation

*Scrib^rumz/rumz^* neural tissue overall exhibits decreased CE, and in particular, decreased narrowing (convergence). *Scrib^rumz/rumz^* mutants exhibit disrupted polarity of cell intercalation, as well as a loss of the normal basal elongation and orientation of neural cells. Interestingly, this phenotype resembles that seen in embryos lacking Ptk7 (*Ptk7^XST87/XST87^)* but not *Vangl2^Lp/Lp^* mutants where decreased CE is due to *decreased frequency* of cell intercalation rather than *defects in polarity* of cell intercalation or basal cell orientation (Williams et al., 2014). The different effects of *Scrib* and *Vangl2^Lp^* mutants on CE and cell intercalation behavior suggest that they are not primarily regulating the same cellular processes during neural morphogenesis, and that the genetic interaction between these mutants may lie in complementary effects on cell behavior. Thus, the effects of Scrib on the *polarity* of intercalation may synergize with those of Vangl2^Lp^ on *frequency* of intercalation.

Furthermore, the similarities between *Scrib^rumz/rumz^* and *Ptk7^XST87/XST87^* embryos suggest that Scrib and Ptk7 may act on similar downstream effectors such as MIIB which is disorganized in *Ptk7^XST87/XST87^* mutant neural tissue or Rho Kinase (ROCK) which regulates cell shape changes needed for NTC in *Ptk7^XST87/XST87^* embryos (Williams et al., 2014; Ybot-Gonzalez et al., 2007). However, Scrib and Ptk7 likely act on these targets through parallel pathways as the *Scrib^Crc^* mutation does not interact genetically with the *Ptk7^chuzhoi^* mutant to cause NTD (Paudyal et al., 2010).

While the *Scrib^rumz^* mutation does not significantly affect the frequency of different types of cell rearrangements, a key observation is the very distinct effect that it has on cell behavior in rosette formations. The loss of just one normal allele not only leads to a randomization of the polarity of rosette formation and junctional resolution, such that rosettes will form from ML and resolve to AP equally as often as they form from AP and resolve to ML, but also leads to rosettes that *form and then resolve back in the same direction* (ML to ML or AP to AP). With the loss of two normal alleles, we observe that any junctional rearrangements of rosettes either lead to resolution back along the original axis, or to repetitive formation and resolution of rosettes of the same groups of cells, without any net change in the position of the cells. In addition, in mutant embryos we also observed numerous rosettes that formed and remained as rosettes for the rest of the movie. Interestingly, we also observed these behaviors in *Vangl2^Lp^* embryos, where the polarity of resolution was randomized, and many rosettes were observed to remain in rosette conformation without resolution (Williams et al., 2014). These results suggest that rosette resolution is critically dependent on Scrib and Vangl2 function.

Comparison of the *Scrib^rumz^*, *Vangl2^Lp^, and Ptk7^Xst87^* phenotypes raises interesting questions about the role of cell intercalation as a force-producing process that actively promotes elongation of the body axis during the early period of neural tube formation in the mouse. All three of these mutants strongly affect either the frequency or polarity of cell intercalation, but only the *Scrib^rumz/rumz^* and *Ptk7^XST87/XST87^* mutants significantly decrease overall CE of the body axis during this period. This suggests that mediolateral cell intercalation may not be the sole, or even the primary driver of neural CE during this period, but may cooperate with other mechanisms. One likely mechanism is an extrinsically generated biomechanical force, either from amniotic fluid pressure (Imuta et al., 2014), or alternatively, from tissue strain generated by the formation and enlargement of the headfolds (Gavrilov and Lacy, 2013; Sutherland et al., 2020). Studies of germband elongation in *Drosophila* have shown that the tensile force generated by posterior midgut invagination drives tissue CE as well as orienting cell rearrangement (Collinet et al., 2015; Lye et al., 2015; Yu and Fernandez-Gonzalez, 2016), and that in the absence of posterior midgut invagination, cell intercalation is not sufficient to drive tissue elongation (Collinet et al., 2015; Yu and Fernandez-Gonzalez, 2016). Accordingly, in the *Vangl2^Lp^* neural plates normal extrinsic forces may be successfully promoting extension despite the changes in cell intercalation. In the *Scrib^rumz/rumz^* and *Ptk7^XST87/XST87^* mutants effects on actin and myosin organization (present data and (Andreeva et al., 2014; Williams et al., 2014)), may be changing the compliance of the tissue and reducing its response to normal extrinsic forces, leading to significantly decreased neural CE.

### Junctional Integrity

Apical constriction and rosette resolution are disrupted in *Scrib^rumz/rumz^* mutant cells, effects which may be due to changes in apical junctional integrity. ZO-1, Par6, Par3, E-cad, and N-cad expression are decreased in the neural plate of *Scrib^rumz/rumz^* mutants. This is consistent with results of Scrib deletion in the head ectoderm of mouse embryos, which led to decreased E-cad and ZO-1 expression and aberrant cell shape changes in the cornea (Yamben et al., 2013), as well as with a model of Scrib regulation of junctional integrity established in Madin Darby Canine Kidney (MDCK) cells (Yamanaka et al., 2003). In this model, loss of Scrib frees up Lethal giant larvae (Lgl), a polarity protein in the basal Scrib-Discs large (Dlg)-Lgl complex, to bind to aPKC and Par6 creating competition between Lgl and Par3 and subsequently inhibiting the formation of tight junctions (Yamanaka et al., 2003). In the neural plate, the *Scrib^rumz^* mutation may disrupt interaction of Lgl with the Scrib LRR domain (Abedrabbo and Ravid, 2020; Kallay et al., 2006) allowing Lgl to localize to the apical membrane and bind to aPKC and Par6. The formation of an Lgl-aPKC-Par6 complex would displace Par3 from the complex and destabilize tight junctions leading to loss of ZO-1 and Par3 expression, which is what we observe in *Scrib^rumz/rumz^* mutants. Additionally, competition with Lgl may displace Par6 from the apical junctions, consistent with our observation that Par6 apical expression is decreased in *Scrib^rumz/rumz^* cells.

Scrib is important for retention and recycling of cadherin at the adherens junctions (Lohia et al., 2012); therefore, loss of Scrib would be expected to result in higher turnover and degradation of cadherin, consistent with our observations of decreased junctional localization of E-cad and N-cad through whole mount staining. How increased turnover of cadherins would result in greater resistance of the tissue to cell rearrangement, including passive rearrangement due to external forces, at first seems contradictory, but the loss of cadherins may result in other adherens junction components, such as nectins, promoting altered cell adhesion dynamics. Alternatively, loss of cadherin-based adhesion could result in increased cell-matrix associations that could be inimicable to passive cell intercalation (Marsden and DeSimone, 2003).

### Actomyosin Pulsing and Biomechanics

Anchoring of the cytoskeleton to the junction is dependent on the presence of cadherins (Jodoin et al., 2015; Martin et al., 2010; Martin and Goldstein, 2014; Sawyer et al., 2011). Whole mount staining revealed that actin and MIIB are decreased while MIIA is increased at neural cell junctions in *Scrib^rumz/rumz^* mutant embryos. The decreased expression of actin and MIIB could be due to disrupted junctional integrity and loss of cadherins, leading to unstable anchorage of the apical cytoskeleton to the junctions. Additionally, it could suggest slower turnover or higher degradation of actin and MIIB at junctions. Increased junctional expression of MIIA was surprising as MIIB and MIIA are often thought to have distinct cellular functions, and typically are not upregulated in response to loss of one or another isoform (Wang et al., 2010). However, MIIA is able to compensate for the loss of MIIB to maintain cell-cell adhesion in neural epithelial tissue (Ma and Adelstein, 2014; Wang et al., 2010). Therefore, it is possible that the increased MIIA in absence of MIIB in the *Scrib^rumz/rumz^* neural plate maintains cellular interactions at cellular junctions, but in a manner that doesn’t promote cell intercalation. MIIA and MIIB are known to have different kinetic profiles (Kovács et al., 2003), and thus the change from MIIB to MIIA might alter cell intercalation behavior while preserving cell-cell adhesion. Alternatively, recent studies in human breast cancer cell lines demonstrated that MIIA co-immunoprecipitated with Lgl but not Scrib, and MIIB complexed with Scrib but not Lgl, demonstrating MIIA and MIIB form distinct complexes with Lgl and Scrib respectively (Abedrabbo and Ravid, 2020). Thus, the increase in MIIA we observe in the neural plate of *Scrib^rumz^* mutants may be linked to increased Lgl at the apical membrane, whereas the decreased expression of MIIB may be directly due to lack of Scrib localization to cell junctions. Future studies using live imaging of actin and myosin in the neural plate will be helpful to elucidate these dynamics.

Not only is the expression of myosin decreased, but the activation of myosin is also decreased in *Scrib^rumz/rumz^* neural tissue. This result is consistent with studies during *Drosophila* germband extension where phospho mutants of the regulatory myosin light chain severely disrupted axial elongation (Kasza et al., 2014). Furthermore, inhibiting myosin activation decreased cell rearrangements during germband extension; while myosin activation promoted rosette rearrangements (Kasza et al., 2014). The defects in productive rosette rearrangements during CE in *Scrib^rumz/rumz^* embryos, as well as defects in expression and activation of actin and myosin are consistent with Scrib playing an essential role in promoting the actomyosin contractility known to be important for CE and apical constriction during embryonic development (Coravos et al., 2017; Heer and Martin, 2017; Murrell et al., 2015; Sutherland and Lesko, 2020). Although it has yet to be demonstrated in mouse neural tube, it is known that actin turnover and stabilization are important for pulsing forces during *Drosophila* embryo development (Dehapiot et al., 2020; Jodoin et al., 2015). Additionally, junctional remodeling is needed to promote a ratchet mechanism for pulsing where contractions promote junctional shrinkage and rearrangement (Rauzi et al., 2010). Our results suggest that turnover and/or junctional integrity may be disrupted in *Scrib^rumz/rumz^* mutants, suggesting that altered actomyosin pulsing is a possible mechanism for the defects in cell shape changed and cell behavior caused by the Scrib mutation. Actomyosin pulsing contractions rely on a Rho-mediated mechanism (Bement et al., 2015; Dehapiot et al., 2020; Garcia De Las Bayonas et al., 2019; Mason et al., 2016; Michaux and Robin, 2018; Munjal et al., 2015; Reyes et al., 2014), which is disrupted in *Ptk7^XST87/XST87^* mutant embryos with neural cell behavior defects that closely resemble the CE and apical constriction phenotypes we observed in *Scrib^rumz/rumz^* mutant embryos (Andreeva et al., 2014; Williams et al., 2014). In MCF7 polarized cysts, Scrib acts as a scaffold to target the RhoGAP protein DCL3 to cell-cell contacts in order to mediate Rho signaling (Hendrick et al., 2016). Therefore, further studies focused on the role of Scrib in regulating Rho-mediated actomyosin pulsing warrant future investigation in order to better understand the genesis of the CE and apical constriction defects observed in *Scrib^rumz/rumz^* mutants.

Not only did we observe CRN in *Scrib^rumz/rumz^* mutant embryos, but we also found prominent turning defects where mutant embryos exhibited axial torsion and, in some cases, a complete lack of turning. Mouse embryos usually turn on axis at around 10-12 somites, however how this process occurs is not fully understood. Turning defects in *Scrib^rumz/rumz^* mutants could be caused by the lack of CE resulting in a wider flatter neural plate as NTC progresses, thereby inhibiting the buildup of forces needed to initiate turning. Another possibility is that disrupted apical constriction in *Scrib^rumz/rumz^* embryos does not allow bending of the neural tube which may alleviate stress on the tissue, which is normally necessary for the embryo to turn. The elevation of the neural folds toward the concave side of the embryonic axis requires increase of tensile elongation forces in the ventral neural/mesodermal tissues or increased compressive forces in the presumptive dorsal (neural fold) region of the neural plate. Finally, junctional composition, cytoskeleton organization, and actomyosin pulsing are required for biomechanical sensing and force propagation (Coravos et al., 2017; Galea et al., 2017; Heer and Martin, 2017; Sutherland and Lesko, 2020; Tharp and Weaver, 2018). Moreover, tissue strain can regulate cell polarity and differentiation at these stages of development (Chien et al., 2015; Chien et al., 2018). We have shown disruption in junctional stability and cytoskeletal organization in *Scrib^rumz/rumz^* neural cells suggesting these processes may be required for the mechanical sensing necessary for embryos to turn. Further studies are needed to determine whether this mechanism is at work in neural morphogenesis.

## Conclusions

Here we investigate the effect of *Scrib* mutation on NTC and determine details of neural cell behavior regulated by Scrib. We show that *Scrib* mutants exhibit CRN and turning defects and genetically interact with *Vangl2^Lp^* mutation to cause NTD. CE, specifically narrowing, is disrupted in *Scrib^rumz/rumz^* mutant embryos, as well as, polarity of cell intercalations and resolution of rosettes as a mechanism of cell rearrangement. Apical constriction and cell shape changes are also disrupted in *Scrib^rumz/rumz^* mutants and we identified a role for Scrib in regulating junctional composition and cytoskeletal organization in the neural plate. Together these studies identify a novel role for Scrib in regulating neural cell behavior and provide a better understanding of NTC in mammalian development

## Acknowledgements

We acknowledge the Keck Center for Cellular Imaging for the usage of the Zeiss 780 microscopy system (PI: Ammasi Periasamy; NIH-ODO16446) and the Leica SP5X microscopy system (PI: Ammasi Periasamy; NIH-RR025616). We also thank Dr. Xiaowei Lu for her helpful discussions and comments.

## Funding Sources

This research was supported by the Eunice Kennedy Shriver National Institute of Child Health and Human Development-R01HD087093.

---

**Supplemental Table 1.**
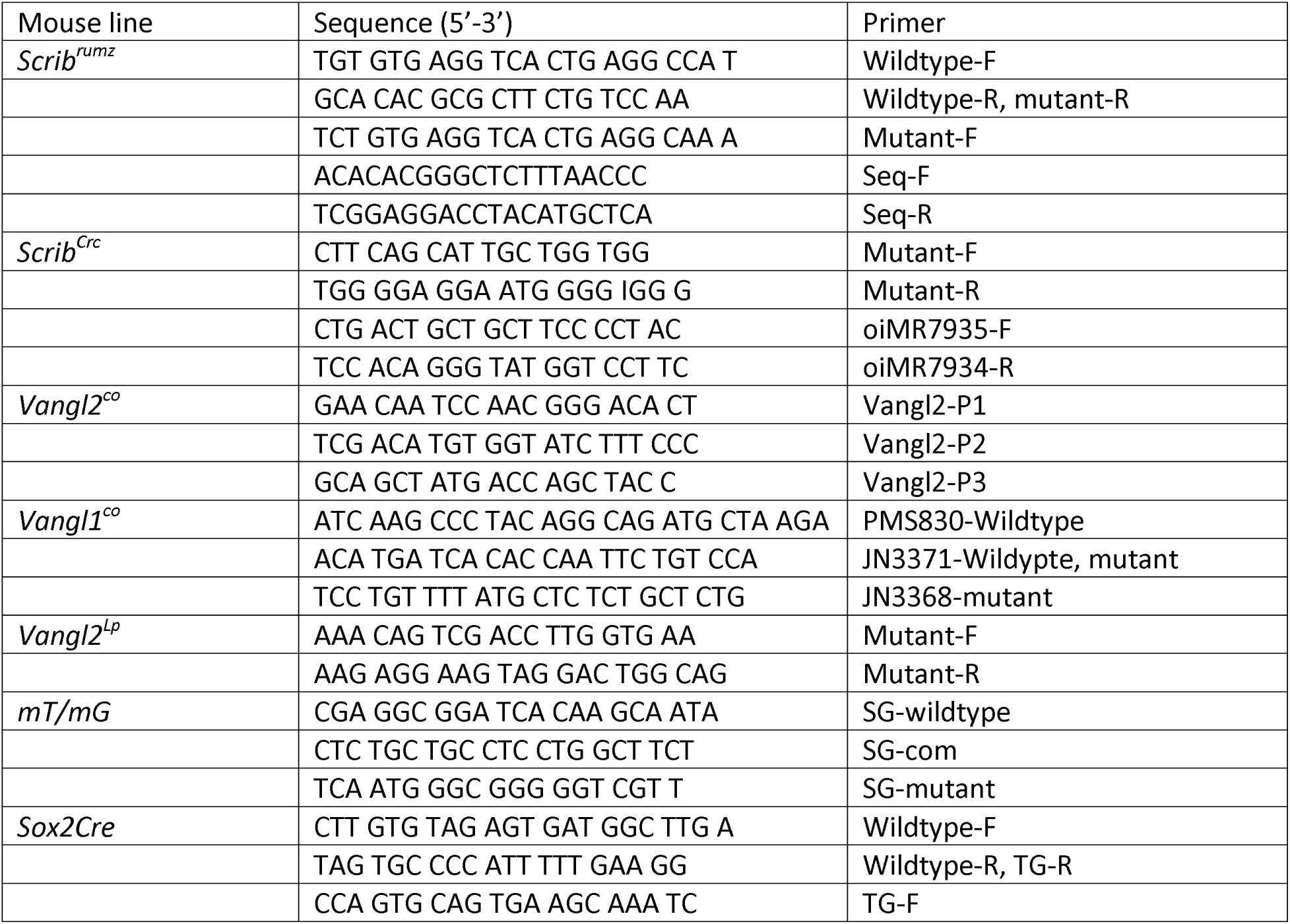
Genotyping Primers. Primer sequences used for genotyping mouse lines by PCR or sequencing.

**Supplemental Table 2.** Characterization of *Scrib* mutant phenotypes. Tables display the percent of embryos exhibiting CRN, exencephaly, turning defects, kinked tail, and developmental delay. Both *Scrib^rumz/rumz^* and *Scrib^Crc/Crc^* mutants exhibit a significantly larger percentage of embryos with CRN and turning defects; however, *Scrib^rumz/rumz^* mutants exhibit a significantly larger percentage of embryos with exencephaly, while *Scrib^Crc/Crc^* mutants exhibit a significantly larger percentage of embryos with kinked tail. Significance is marked with * and was determined within each phenotype by Fishers Exact test compared to wildtype littermates. n represents the number of embryos characterized.

|  | Wildtype<br>(n=27) | *Scrib*<sup>rumz/+</sup><br>(n=58) | *Scrib*<sup>rumz/rumz</sup><br>(n=26) |
| --- | --- | --- | --- |
| craniorachischisis | 0.0 | 3.6<br>p= 0.1212 | 80.8*<br>p= <0.0001 |
| exencephaly | 0.0 | 3.4<br>p= 0.2462 | 7.7*<br>p= 0.0068 |
| turning defect | 0.0 | 3.4<br>p= 0.2462 | 73.1*<br>p= <0.0001 |
| kink tail | 14.8 | 24.1<br>p= 0.1528 | 15.4<br>p=>0.9999 |
| developmental delay | 7.4 | 12.1<br>p= 0.3350 | 15.4<br>p= 0.1121 |

|  | Wildtype<br>(n=30) | *Scrib*<sup>Crc/+</sup><br>(n=49) | *Scrib*<sup>Crc/Crc</sup><br>(n=26) |
| --- | --- | --- | --- |
| craniorachischisis | 0.0 | 2.0<br>p=0.4975 | 88.5*<br>p=<0.0001 |
| exencephaly | 0.0 | 4.14<br>p=0.1212 | 3.8<br>p=0.1212 |
| turning defect | 0.0 | 4.1<br>p=0.1212 | 53.8*<br>p=<0.0001 |
| kink tail | 16.7 | 18.4<br>p=>0.9999 | 46.2*<br>p=<0.0001 |
| developmental delay | 3.3 | 6.1<br>p=0.4977 | 7.7<br>p=0.2134 |
| Note: While both *Scrib*<sup>rumz/rumz</sup> and *Scrib*<sup>Crc/Crc</sup> have prominent defects in CRN and turning, exencephaly is significantly increased in *Scrib*<sup>rumz/rumz</sup> mutants and kinked tail is significantly increased in *Scrib*<sup>Crc/Crc</sup> mutants. |  |  |  |

**Supplemental Figure 1.**
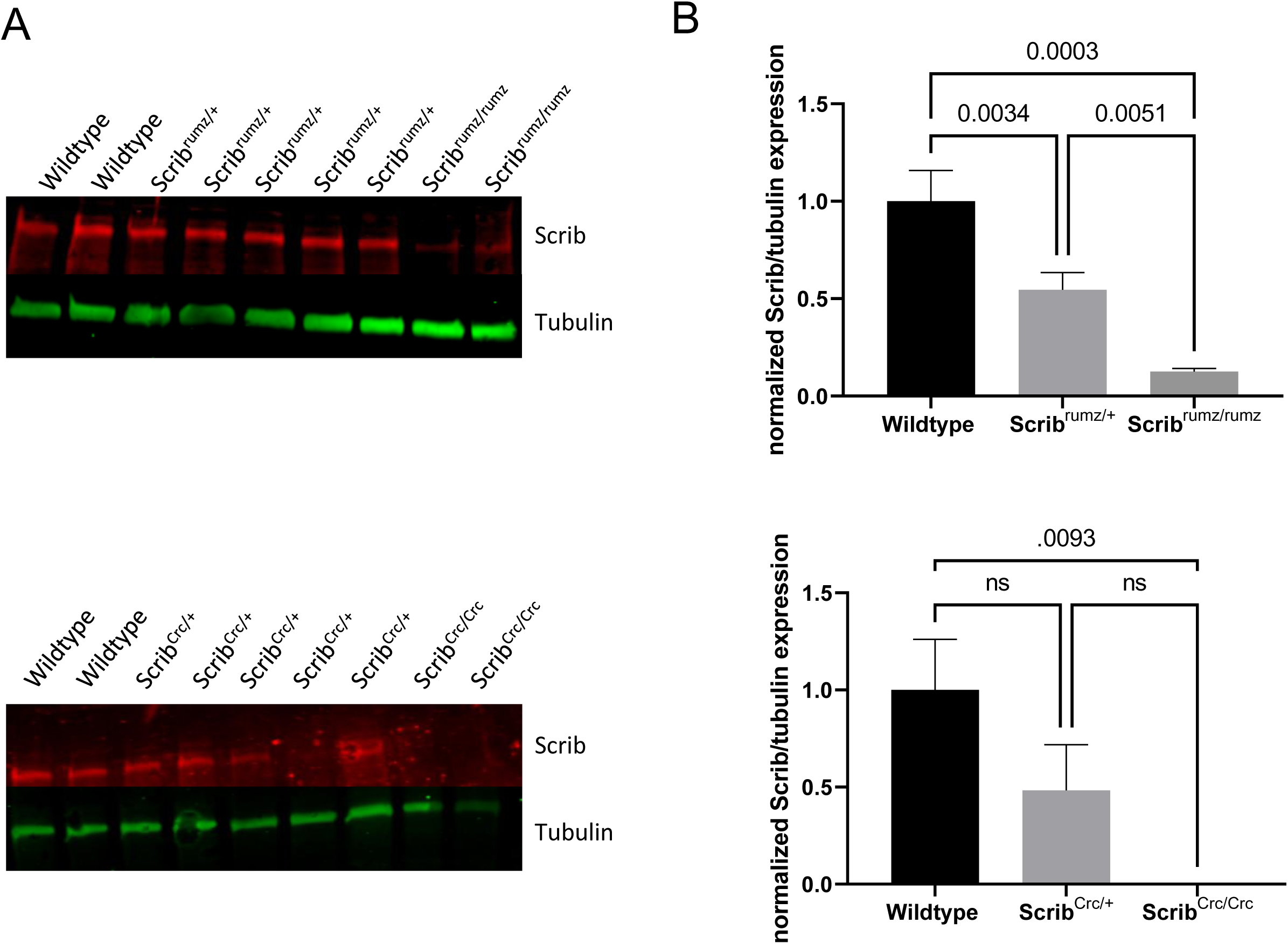
Protein expression in *Scrib* mutants. A) Representative western blots for Scrib (red) and tubulin (green) in *Scrib^rumz^* (top) and *Scrib^Crc^* (bottom) embryos. B) Western blots were analyzed with Image Studio and normalized to tubulin expression. Error bars represent standard deviation and p-values were determined by one-way ANOVA (p<0.05) and ns denotes not significant (p>0.05).

**Supplemental Figure 2.**
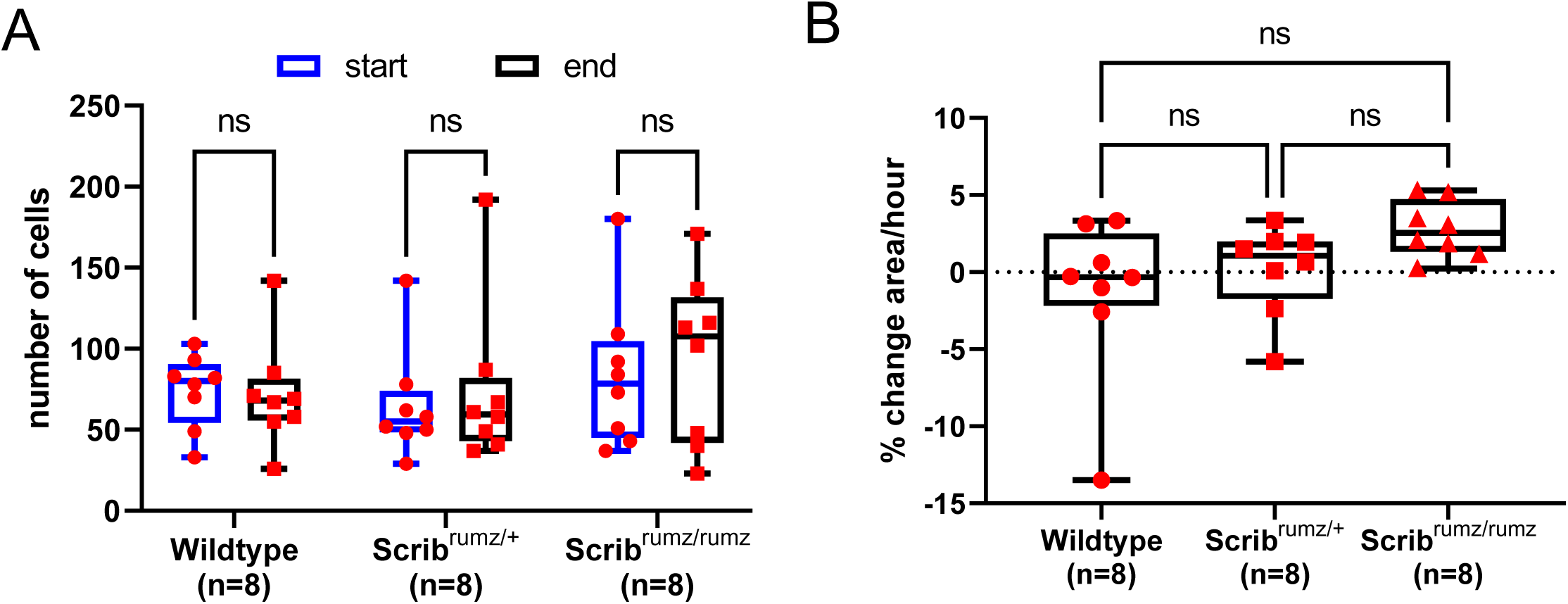
Overall tissue shape changes in the neural tube is due to cell shape changes and not cell number. A) Graph indicates number of cells within distortion diagrams at the start and end of live confocal movies and shows no counts are statistically significant from one another determined by two-way ANOVA (ns denotes not significant, p>0.05). n represents number of embryos analyzed. Error bars represent min and max. B) Plot of percent change in area of distortion diagrams per hour. n represents the number of movies analyzed. Error bars represent min and max. ns denotes not significant (p>0.05) and was determined by two-way ANOVA.

## Notes

### Competing Interest Statement

The authors have declared no competing interest.

## Reference

Abedrabbo, M., Ravid, S., 2020. Scribble, Lgl1, and myosin II form a complex in vivo to promote directed cell migration. Molecular biology of the cell, mbcE19110657.

Andreeva, A., Lee, J., Lohia, M., Wu, X., Macara, I.G., Lu, X., 2014. PTK7-Src signaling at epithelial cell contacts mediates spatial organization of actomyosin and planar cell polarity. Developmental cell 29, 20–33.

Axelrod, J.D., McNeill, H., 2002. Coupling planar cell polarity signaling to morphogenesis. TheScientificWorldJournal 2, 434–454.

Belotti, E., Polanowska, J., Daulat, A.M., Audebert, S., Thomé, V., Lissitzky, J.C., Lembo, F., Blibek, K., Omi, S., Lenfant, N., Gangar, A., Montcouquiol, M., Santoni, M.J., Sebbagh, M., Aurrand-Lions, M., Angers, S., Kodjabachian, L., Reboul, J., Borg, J.P., 2013. The human PDZome: a gateway to PSD95-Disc large-zonula occludens (PDZ)-mediated functions. Molecular & cellular proteomics : MCP 12, 2587–2603.

Bement, W.M., Leda, M., Moe, A.M., Kita, A.M., Larson, M.E., Golding, A.E., Pfeuti, C., Su, K.C., Miller, A.L., Goryachev, A.B., von Dassow, G., 2015. Activator-inhibitor coupling between Rho signalling and actin assembly makes the cell cortex an excitable medium. Nature cell biology 17, 1471–1483.

Bertet, C., Sulak, L., Lecuit, T., 2004. Myosin-dependent junction remodelling controls planar cell intercalation and axis elongation. Nature 429, 667–671.

Blankenship, J.T., Backovic, S.T., Sanny, J.S., Weitz, O., Zallen, J.A., 2006. Multicellular rosette formation links planar cell polarity to tissue morphogenesis. Developmental cell 11, 459–470.

Blencowe, H., Kancherla, V., Moorthie, S., Darlison, M.W., Modell, B., 2018. Estimates of global and regional prevalence of neural tube defects for 2015: a systematic analysis. Annals of the New York Academy of Sciences 1414, 31–46.

Bonello, T.T., Peifer, M., 2018. Scribble: A master scaffold in polarity, adhesion, synaptogenesis, and proliferation.

Butler, M.B., Short, N.E., Maniou, E., Alexandre, P., Greene, N.D.E., Copp, A.J., Galea, G.L., 2019. Rho kinase-dependent apical constriction counteracts M-phase apical expansion to enable mouse neural tube closure. 132.

Chang, H., Smallwood, P.M., Williams, J., Nathans, J., 2016. The spatio-temporal domains of Frizzled6 action in planar polarity control of hair follicle orientation. Developmental biology 409, 181–193.

Chien, Y.H., Keller, R., Kintner, C., Shook, D.R., 2015. Mechanical strain determines the axis of planar polarity in ciliated epithelia. Current biology : CB 25, 2774–2784.

Chien, Y.H., Srinivasan, S., Keller, R., Kintner, C., 2018. Mechanical Strain Determines Cilia Length, Motility, and Planar Position in the Left-Right Organizer. Developmental cell 45, 316–330.e314.

Collinet, C., Rauzi, M., Lenne, P.F., Lecuit, T., 2015. Local and tissue-scale forces drive oriented junction growth during tissue extension. Nature cell biology 17, 1247–1258.

Copp, A.J., Greene, N.D., 2010. Genetics and development of neural tube defects. The Journal of pathology 220, 217–230.

Copp, A.J., Greene, N.D., 2013. Neural tube defects--disorders of neurulation and related embryonic processes. Wiley interdisciplinary reviews. Developmental biology 2, 213–227.

Coravos, J.S., Mason, F.M., Martin, A.C., 2017. Actomyosin Pulsing in Tissue Integrity Maintenance during Morphogenesis. Trends in cell biology 27, 276–283.

Courbard, J.R., Djiane, A., Wu, J., Mlodzik, M., 2009. The apical/basal-polarity determinant Scribble cooperates with the PCP core factor Stbm/Vang and functions as one of its effectors. Developmental biology 333, 67–77.

Dehapiot, B., Clément, R., Alégot, H., Gazsó-Gerhát, G., Philippe, J.M., Lecuit, T., 2020. Assembly of a persistent apical actin network by the formin Frl/Fmnl tunes epithelial cell deformability. Nature cell biology.

Dow, L.E., Kauffman, J.S., Caddy, J., Zarbalis, K., Peterson, A.S., Jane, S.M., Russell, S.M., Humbert, P.O., 2007. The tumour-suppressor Scribble dictates cell polarity during directed epithelial migration: regulation of Rho GTPase recruitment to the leading edge. Oncogene 26, 2272–2282.

Escuin, S., Vernay, B., Savery, D., Gurniak, C.B., Witke, W., Greene, N.D., Copp, A.J., 2015. Rho-kinase-dependent actin turnover and actomyosin disassembly are necessary for mouse spinal neural tube closure. Journal of cell science 128, 2468–2481.

Fernandez-Gonzalez, R., Zallen, J.A., 2011. Oscillatory behaviors and hierarchical assembly of contractile structures in intercalating cells. Physical biology 8, 045005.

Fristrom, D., 1976. The mechanism of evagination of imaginal discs of Drosophila melanogaster. III. Evidence for cell rearrangement. Developmental biology 54, 163–171.

Galea, G.L., Cho, Y.J., Galea, G., Mole, M.A., Rolo, A., 2017. Biomechanical coupling facilitates spinal neural tube closure in mouse embryos. 114, E5177–e5186.

Galea, G.L., Nychyk, O., Mole, M.A., Moulding, D., Savery, D., Nikolopoulou, E., Henderson, D.J., Greene, N.D.E., Copp, A.J., 2018. Vangl2 disruption alters the biomechanics of late spinal neurulation leading to spina bifida in mouse embryos. Disease models & mechanisms 11.

Garcia De Las Bayonas, A., Philippe, J.M., Lellouch, A.C., Lecuit, T., 2019. Distinct RhoGEFs Activate Apical and Junctional Contractility under Control of G Proteins during Epithelial Morphogenesis. Current biology: CB.

Gavrilov, S., Lacy, E., 2013. Genetic dissection of ventral folding morphogenesis in mouse: embryonic visceral endoderm-supplied BMP2 positions head and heart. Current opinion in genetics & development 23, 461–469.

Gravel, M., Iliescu, A., Horth, C., Apuzzo, S., Gros, P., 2010. Molecular and cellular mechanisms underlying neural tube defects in the loop-tail mutant mouse. Biochemistry 49, 3445–3455.

Greene, N.D., Copp, A.J., 2014. Neural tube defects. Annual review of neuroscience 37, 221–242.

Hamblet, N.S., Lijam, N., Ruiz-Lozano, P., Wang, J., Yang, Y., Luo, Z., Mei, L., Chien, K.R., Sussman, D.J., Wynshaw-Boris, A., 2002. Dishevelled 2 is essential for cardiac outflow tract development, somite segmentation and neural tube closure. Development (Cambridge, England) 129, 5827–5838.

Hayashi, S., Tenzen, T., McMahon, A.P., 2003. Maternal inheritance of Cre activity in a Sox2Cre deleter strain. Genesis (New York, N.Y.: 2000) 37, 51–53.

Heer, N.C., Martin, A.C., 2017. Tension, contraction and tissue morphogenesis. 144, 4249–4260.

Heisenberg, C.P., Tada, M., Rauch, G.J., Saude, L., Concha, M.L., Geisler, R., Stemple, D.L., Smith, J.C., Wilson, S.W., 2000. Silberblick/Wnt11 mediates convergent extension movements during zebrafish gastrulation. Nature 405, 76–81.

Hendrick, J., Franz-Wachtel, M., Moeller, Y., Schmid, S., Macek, B., Olayioye, M.A., 2016. The polarity protein Scribble positions DLC3 at adherens junctions to regulate Rho signaling. Journal of cell science 129, 3583–3596.

Iliescu, A., Gravel, M., Horth, C., Apuzzo, S., Gros, P., 2011. Transmembrane topology of mammalian planar cell polarity protein Vangl1. Biochemistry 50, 2274–2282.

Imuta, Y., Koyama, H., Shi, D., Eiraku, M., Fujimori, T., Sasaki, H., 2014. Mechanical control of notochord morphogenesis by extra-embryonic tissues in mouse embryos. Mechanisms of development 132, 44–58.

Inoue, Y., Suzuki, M., Watanabe, T., Yasue, N., Tateo, I., Adachi, T., Ueno, N., 2016. Mechanical roles of apical constriction, cell elongation, and cell migration during neural tube formation in Xenopus. Biomechanics and modeling in mechanobiology 15, 1733–1746.

Irvine, K.D., Wieschaus, E., 1994. Cell intercalation during Drosophila germband extension and its regulation by pair-rule segmentation genes. Development (Cambridge, England) 120, 827–841.

Jodoin, J.N., Coravos, J.S., Chanet, S., Vasquez, C.G., Tworoger, M., Kingston, E.R., Perkins, L.A., Perrimon, N., Martin, A.C., 2015. Stable Force Balance between Epithelial Cells Arises from F-Actin Turnover. Developmental cell 35, 685–697.

Kallay, L.M., McNickle, A., Brennwald, P.J., Hubbard, A.L., Braiterman, L.T., 2006. Scribble associates with two polarity proteins, Lgl2 and Vangl2, via distinct molecular domains. Journal of cellular biochemistry 99, 647–664.

Kasza, K.E., Farrell, D.L., Zallen, J.A., 2014. Spatiotemporal control of epithelial remodeling by regulated myosin phosphorylation. Proceedings of the National Academy of Sciences of the United States of America 111, 11732–11737.

Keller, R., 2002. Shaping the vertebrate body plan by polarized embryonic cell movements. Science (New York, N.Y.) 298, 1950–1954.

Keller, R., Davidson, L., Edlund, A., Elul, T., Ezin, M., Shook, D., Skoglund, P., 2000. Mechanisms of convergence and extension by cell intercalation. Philosophical transactions of the Royal Society of London. Series B, Biological sciences 355, 897–922.

Keller, R., Sutherland, A., 2020. Convergent extension in the amphibian, Xenopus laevis. Current topics in developmental biology 136, 271–317.

Keller, R.E., 1978. Time-lapse cinemicrographic analysis of superficial cell behavior during and prior to gastrulation in Xenopus laevis. Journal of morphology 157, 223–247.

Kharfallah, F., Guyot, M.C., El Hassan, A.R., Allache, R., Merello, E., De Marco, P., Di Cristo, G., Capra, V., Kibar, Z., 2017. Scribble1 plays an important role in the pathogenesis of neural tube defects through its mediating effect of Par-3 and Vangl1/2 localization. Human molecular genetics 26, 2307–2320.

Kibar, Z., Vogan, K.J., Groulx, N., Justice, M.J., Underhill, D.A., Gros, P., 2001. Ltap, a mammalian homolog of Drosophila Strabismus/Van Gogh, is altered in the mouse neural tube mutant Loop-tail. Nature genetics 28, 251–255.

Kinoshita, N., Sasai, N., Misaki, K., Yonemura, S., 2008. Apical accumulation of Rho in the neural plate is important for neural plate cell shape change and neural tube formation. Molecular biology of the cell 19, 2289–2299.

Kovács, M., Wang, F., Hu, A., Zhang, Y., Sellers, J.R., 2003. Functional divergence of human cytoplasmic myosin II: kinetic characterization of the non-muscle IIA isoform. The Journal of biological chemistry 278, 38132–38140.

Lei, Y., Zhu, H., Duhon, C., Yang, W., Ross, M.E., Shaw, G.M., Finnell, R.H., 2013. Mutations in planar cell polarity gene SCRIB are associated with spina bifida. PloS one 8, e69262.

Lohia, M., Qin, Y., Macara, I.G., 2012. The Scribble polarity protein stabilizes E-cadherin/p120-catenin binding and blocks retrieval of E-cadherin to the Golgi. PloS one 7, e51130.

López-Escobar, B., Caro-Vega, J.M., Vijayraghavan, D.S., Plageman, T.F., Sanchez-Alcazar, J.A., Moreno, R.C., Savery, D., Márquez-Rivas, J., Davidson, L.A., Ybot-González, P., 2018. The non-canonical Wnt-PCP pathway shapes the mouse caudal neural plate. Development (Cambridge, England) 145.

Lye, C.M., Blanchard, G.B., Naylor, H.W., Muresan, L., Huisken, J., Adams, R.J., Sanson, B., 2015. Mechanical Coupling between Endoderm Invagination and Axis Extension in Drosophila. PLoS biology 13, e1002292.

Ma, X., Adelstein, R.S., 2014. The role of vertebrate nonmuscle Myosin II in development and human disease. Bioarchitecture 4, 88–102.

Marsden, M., DeSimone, D.W., 2003. Integrin-ECM interactions regulate cadherin-dependent cell adhesion and are required for convergent extension in Xenopus. Current biology : CB 13, 1182–1191.

Martin, A.C., Gelbart, M., Fernandez-Gonzalez, R., Kaschube, M., Wieschaus, E.F., 2010. Integration of contractile forces during tissue invagination. The Journal of cell biology 188, 735–749.

Martin, A.C., Goldstein, B., 2014. Apical constriction: themes and variations on a cellular mechanism driving morphogenesis. Development (Cambridge, England) 141, 1987–1998.

Mason, F.M., Xie, S., Vasquez, C.G., Tworoger, M., Martin, A.C., 2016. RhoA GTPase inhibition organizes contraction during epithelial morphogenesis. The Journal of cell biology 214, 603–617.

McShane, S.G., Mole, M.A., Savery, D., Greene, N.D., Tam, P.P., Copp, A.J., 2015. Cellular basis of neuroepithelial bending during mouse spinal neural tube closure. Developmental biology 404, 113–124.

Merte, J., Jensen, D., Wright, K., Sarsfield, S., Wang, Y., Schekman, R., Ginty, D.D., 2010. Sec24b selectively sorts Vangl2 to regulate planar cell polarity during neural tube closure. Nature cell biology 12, 41–46; sup pp 41-48.

Michaux, J.B., Robin, F.B., 2018. Excitable RhoA dynamics drive pulsed contractions in the early C. elegans embryo. 217, 4230–4252.

Montcouquiol, M., Sans, N., Huss, D., Kach, J., Dickman, J.D., Forge, A., Rachel, R.A., Copeland, N.G., Jenkins, N.A., Bogani, D., Murdoch, J., Warchol, M.E., Wenthold, R.J., Kelley, M.W., 2006. Asymmetric localization of Vangl2 and Fz3 indicate novel mechanisms for planar cell polarity in mammals. The Journal of neuroscience : the official journal of the Society for Neuroscience 26, 5265–5275.

Moury, J.D., Schoenwolf, G.C., 1995. Cooperative model of epithelial shaping and bending during avian neurulation: autonomous movements of the neural plate, autonomous movements of the epidermis, and interactions in the neural plate/epidermis transition zone. Developmental dynamics : an official publication of the American Association of Anatomists 204, 323–337.

Munjal, A., Philippe, J.M., Munro, E., Lecuit, T., 2015. A self-organized biomechanical network drives shape changes during tissue morphogenesis. Nature 524, 351–355.

Munro, E.M., Odell, G.M., 2002. Polarized basolateral cell motility underlies invagination and convergent extension of the ascidian notochord. Development (Cambridge, England) 129, 13–24.

Murdoch, J.N., Damrau, C., Paudyal, A., Bogani, D., Wells, S., Greene, N.D., Stanier, P., Copp, A.J., 2014. Genetic interactions between planar cell polarity genes cause diverse neural tube defects in mice. Disease models & mechanisms 7, 1153–1163.

Murdoch, J.N., Doudney, K., Paternotte, C., Copp, A.J., Stanier, P., 2001a. Severe neural tube defects in the loop-tail mouse result from mutation of Lpp1, a novel gene involved in floor plate specification. Human molecular genetics 10, 2593–2601.

Murdoch, J.N., Henderson, D.J., Doudney, K., Gaston-Massuet, C., Phillips, H.M., Paternotte, C., Arkell, R., Stanier, P., Copp, A.J., 2003. Disruption of scribble (Scrb1) causes severe neural tube defects in the circletail mouse. Human molecular genetics 12, 87–98.

Murdoch, J.N., Rachel, R.A., Shah, S., Beermann, F., Stanier, P., Mason, C.A., Copp, A.J., 2001b. Circletail, a new mouse mutant with severe neural tube defects: chromosomal localization and interaction with the loop-tail mutation. Genomics 78, 55–63.

Murrell, M., Oakes, P.W., Lenz, M., Gardel, M.L., 2015. Forcing cells into shape: the mechanics of actomyosin contractility. Nature reviews. Molecular cell biology 16, 486–498.

Muzumdar, M.D., Tasic, B., Miyamichi, K., Li, L., Luo, L., 2007. A global double-fluorescent Cre reporter mouse. Genesis 45, 593–605.

Nikolopoulou, E., Galea, G.L., Rolo, A., Greene, N.D., Copp, A.J., 2017. Neural tube closure: cellular, molecular and biomechanical mechanisms. Development (Cambridge, England) 144, 552–566.

Osmani, N., Vitale, N., Borg, J.P., Etienne-Manneville, S., 2006. Scrib controls Cdc42 localization and activity to promote cell polarization during astrocyte migration. Current biology : CB 16, 2395–2405.

Paudyal, A., Damrau, C., Patterson, V.L., Ermakov, A., Formstone, C., Lalanne, Z., Wells, S., Lu, X., Norris, D.P., Dean, C.H., Henderson, D.J., Murdoch, J.N., 2010. The novel mouse mutant, chuzhoi, has disruption of Ptk7 protein and exhibits defects in neural tube, heart and lung development and abnormal planar cell polarity in the ear. BMC developmental biology 10, 87.

Phua, D.C., Humbert, P.O., Hunziker, W., 2009. Vimentin regulates scribble activity by protecting it from proteasomal degradation. Molecular biology of the cell 20, 2841–2855.

Qian, D., Jones, C., Rzadzinska, A., Mark, S., Zhang, X., Steel, K.P., Dai, X., Chen, P., 2007. Wnt5a functions in planar cell polarity regulation in mice. Developmental biology 306, 121–133.

Raman, R., Pinto, C.S., Sonawane, M., 2018. Polarized Organization of the Cytoskeleton: Regulation by Cell Polarity Proteins. Journal of molecular biology 430, 3565–3584.

Rauzi, M., Lenne, P.F., Lecuit, T., 2010. Planar polarized actomyosin contractile flows control epithelial junction remodelling. Nature 468, 1110–1114.

Reyes, C.C., Jin, M., Breznau, E.B., Espino, R., Delgado-Gonzalo, R., Goryachev, A.B., Miller, A.L., 2014. Anillin regulates cell-cell junction integrity by organizing junctional accumulation of Rho-GTP and actomyosin. Current biology : CB 24, 1263–1270.

Robinson, A., Escuin, S., Doudney, K., Vekemans, M., Stevenson, R.E., Greene, N.D., Copp, A.J., Stanier, P., 2012. Mutations in the planar cell polarity genes CELSR1 and SCRIB are associated with the severe neural tube defect craniorachischisis. Human mutation 33, 440–447.

Sadler, T.W., 2005. Embryology of neural tube development. American journal of medical genetics. Part C, Seminars in medical genetics 135c, 2–8.

Sawyer, J.K., Choi, W., Jung, K.C., He, L., Harris, N.J., Peifer, M., 2011. A contractile actomyosin network linked to adherens junctions by Canoe/afadin helps drive convergent extension. Molecular biology of the cell 22, 2491–2508.

Shih, J., Keller, R., 1992. Cell motility driving mediolateral intercalation in explants of Xenopus laevis. Development (Cambridge, England) 116, 901–914.

Shindo, A., 2018. Models of convergent extension during morphogenesis. Wiley interdisciplinary reviews. Developmental biology 7.

Smith, J.L., Schoenwolf, G.C., Quan, J., 1994. Quantitative analyses of neuroepithelial cell shapes during bending of the mouse neural plate. The Journal of comparative neurology 342, 144–151.

Song, H., Hu, J., Chen, W., Elliott, G., Andre, P., Gao, B., Yang, Y., 2010. Planar cell polarity breaks bilateral symmetry by controlling ciliary positioning. Nature 466, 378–382.

Sun, Z., Amourda, C., 2017. Basolateral protrusion and apical contraction cooperatively drive Drosophila germ-band extension. 19, 375–383.

Sutherland, A., Keller, R., Lesko, A., 2020. Convergent extension in mammalian morphogenesis. Seminars in cell & developmental biology 100, 199–211.

Sutherland, A., Lesko, A., 2020. Pulsed actomyosin contractions in morphogenesis. F1000 Research 9.

Suzuki, M., Morita, H., Ueno, N., 2012. Molecular mechanisms of cell shape changes that contribute to vertebrate neural tube closure. Development, growth & differentiation 54, 266–276.

Tada, M., Smith, J.C., 2000. Xwnt11 is a target of Xenopus Brachyury: regulation of gastrulation movements via Dishevelled, but not through the canonical Wnt pathway. Development (Cambridge, England) 127, 2227–2238.

Taylor, J., Adler, P.N., 2008. Cell rearrangement and cell division during the tissue level morphogenesis of evaginating Drosophila imaginal discs. Developmental biology 313, 739–751.

Tharp, K.M., Weaver, V.M., 2018. Modeling Tissue Polarity in Context. Journal of molecular biology 430, 3613–3628.

Torban, E., Wang, H.J., Groulx, N., Gros, P., 2004. Independent mutations in mouse Vangl2 that cause neural tube defects in looptail mice impair interaction with members of the Dishevelled family. The Journal of biological chemistry 279, 52703–52713.

Wallingford, J.B., Harland, R.M., 2001. Xenopus Dishevelled signaling regulates both neural and mesodermal convergent extension: parallel forces elongating the body axis. Development (Cambridge, England) 128, 2581–2592.

Wallingford, J.B., Rowning, B.A., Vogeli, K.M., Rothbacher, U., Fraser, S.E., Harland, R.M., 2000. Dishevelled controls cell polarity during Xenopus gastrulation. Nature 405, 81–85.

Wang, A., Ma, X., Conti, M.A., Liu, C., Kawamoto, S., Adelstein, R.S., 2010. Nonmuscle myosin II isoform and domain specificity during early mouse development. Proceedings of the National Academy of Sciences of the United States of America 107, 14645–14650.

Wang, J., Hamblet, N.S., Mark, S., Dickinson, M.E., Brinkman, B.C., Segil, N., Fraser, S.E., Chen, P., Wallingford, J.B., Wynshaw-Boris, A., 2006a. Dishevelled genes mediate a conserved mammalian PCP pathway to regulate convergent extension during neurulation. Development (Cambridge, England) 133, 1767–1778.

Wang, Y., Guo, N., Nathans, J., 2006b. The role of Frizzled3 and Frizzled6 in neural tube closure and in the planar polarity of inner-ear sensory hair cells. The Journal of neuroscience : the official journal of the Society for Neuroscience 26, 2147–2156.

Williams-Masson, E.M., Heid, P.J., Lavin, C.A., Hardin, J., 1998. The cellular mechanism of epithelial rearrangement during morphogenesis of the Caenorhabditis elegans dorsal hypodermis. Developmental biology 204, 263–276.

Williams, M., Yen, W., Lu, X., Sutherland, A., 2014. Distinct apical and basolateral mechanisms drive planar cell polarity-dependent convergent extension of the mouse neural plate. Developmental cell 29, 34–46.

Wu, G., Huang, X., Hua, Y., Mu, D., 2011. Roles of planar cell polarity pathways in the development of neural [correction of neutral] tube defects. Journal of biomedical science 18, 66.

Yamanaka, T., Horikoshi, Y., Sugiyama, Y., Ishiyama, C., Suzuki, A., Hirose, T., Iwamatsu, A., Shinohara, A., Ohno, S., 2003. Mammalian Lgl forms a protein complex with PAR-6 and aPKC independently of PAR-3 to regulate epithelial cell polarity. Current biology : CB 13, 734–743.

Yamben, I.F., Rachel, R.A., Shatadal, S., Copeland, N.G., Jenkins, N.A., Warming, S., Griep, A.E., 2013. Scrib is required for epithelial cell identity and prevents epithelial to mesenchymal transition in the mouse. Developmental biology 384, 41–52.

Ybot-Gonzalez, P., Savery, D., Gerrelli, D., Signore, M., Mitchell, C.E., Faux, C.H., Greene, N.D., Copp, A.J., 2007. Convergent extension, planar-cell-polarity signalling and initiation of mouse neural tube closure. Development (Cambridge, England) 134, 789–799.

Yen, W.W., Williams, M., Periasamy, A., Conaway, M., Burdsal, C., Keller, R., Lu, X., Sutherland, A., 2009. PTK7 is essential for polarized cell motility and convergent extension during mouse gastrulation. Development (Cambridge, England) 136, 2039–2048.

Yin, H., Copley, C.O., Goodrich, L.V., Deans, M.R., 2012. Comparison of phenotypes between different vangl2 mutants demonstrates dominant effects of the Looptail mutation during hair cell development. PloS one 7, e31988.

Yu, J.C., Fernandez-Gonzalez, R., 2016. Local mechanical forces promote polarized junctional assembly and axis elongation in Drosophila. 5.

Zarbalis, K., May, S.R., Shen, Y., Ekker, M., Rubenstein, J.L., Peterson, A.S., 2004. A focused and efficient genetic screening strategy in the mouse: identification of mutations that disrupt cortical development. PLoS biology 2, E219.

Zohn, I.E., 2020. Mouse Models of Neural Tube Defects. Advances in experimental medicine and biology 1236, 39–64.

